# Effect of Mutations on Smlt1473 Binding to Various Substrates Using Molecular Dynamics Simulations

**DOI:** 10.1101/2024.09.24.614745

**Authors:** Kinjal Mondal, Samantha Felton, Bryan W. Berger, Jeffery B. Klauda

**Affiliations:** Institute for Physical Science and Technology, Biophysics Program, College Park, MD 20742, USA; Department of Biomedical Engineering, University of Virginia, Charlottesville, VA, USA; Department of Chemical Engineering, University of Virginia, Charlottesville, VA, USA; Department of Chemical and Biomolecular Engineering, University of Maryland, College Park, MD 20742, USA

## Abstract

Smlt1473 is a polysaccharide lyase from *Stenotrophomonas maltophilia* whose crystal structure was solved recently using X-ray crystallography. There was an effort to study the effect of mutations on the activity of Smlt1473 binding to various substrates like hyaluronic acid(HA), mannuronic acid(ManA), and alginate. In this study, we use molecular docking and molecular dynamics simulations to investigate the effect of binding of various substrates (HA and ManA) to Smlt1473 and two of its mutants H221F and R312L. We further studied the stability in the binding of Smlt1473 to its various substrates as well as the role of fluctuations. Machine-Learning based clustering algorithms were used to group the entire simulation trajectory into various stable states. The molecular interactions Smlt1473 to the substrates were calculated and the importance of specific residues were tested with observed activity assays due to residue mutations. Overall, we find that the R218 plays an important role in substrate binding and thus impacting the activity due to the H221F mutant and R/L312 itself plays an important role in the R312 mutation. In addition, we have also found three more residues K56, R107, and R164 important for substrate binding which we further proceed to confirm using wet lab mutagenesis studies.

## 1 Introduction

Plants produce complex carbohydrates like starch and cellulose via various biophysical processes like photosynthesis. However these complex carbohydrates must be broken down into simpler sugars before single cell organisms produce chemicals, value-added materials and biofuels^1^. Polysaccharide lyases which are a class of carbohydrate-enzymes that break down long chain anionic polysaccharides through a beta-elimination mechanism can provide a possible solution.

Polysaccharide lyases can be subdivided into 41 subclasses of enzymes based on sequence similarity. This huge diversity has gained sufficient importance in the past decade due to their applications in breaking down a variety of polysaccharides with an even larger diversity in the linkages. Amongst polysaccharide lyases, the alginate lyase is the most well studied with multiple studies related to therapeutics of chronic infection as well as the degradation of multiple poly-mannuronic-acids(M) or poly-glucuronic-acid(G) building blocks ^2–6^.

Some recent studies on polysaccharide lyases have focused on the PL-5 family, including Smlt1473 present in *S. maltophilia*. Smlt1473 was initially shown to have activity towards a variety of polysaccharides such as hyaluronic acid and alginate. Early studies focused on the effect of pH on the activity of Smlt1473 towards its substrates. Broadly, the following pH dependencies were determined for Smlt1473 activity against specific substrates; hyaluronic acid (HA) at pH=5, glucuronic acid (GlcA) at pH=7, and mannuronic acid (ManA) at pH=9^7^. Next a docking and simulation study was conducted in which the interactions of various residues in the enzyme with the sugars were determined ^8^. However, at this point the X-ray crystal structure of Smlt1473 was not determined and the starting structure of Smlt1473 was homology modelled^9^. To increase accuracy of the Smlt1473 structure used in these modeling studies, a subsequent study determined the crystal structures of both the apo as well as the substrate bound Smlt1473 at various pHs using X-ray crystallography. From this study, various residues important to the substrate binding mode of Smlt1473 were identified. Point mutations were made at these different residues to identify potential increase or decrease in activity to give further insight on the interactions of the various residues with the sugar units^10^.

In our study, we aim to use molecular dynamics (MD) simulations to analyze the effects of mutations on the structural changes and predominant effects to gain additional insights into these important binding sites of Smlt1473. We choose to study the mutants H221F and R312L as they display drastic changes in activity compared to the wildtype (WT)^11^. Surprisingly, H221F shows opposing effects on HA as compared to ManA. For HA, H221F drastically decreases activity with respect to the WT, while for ManA, the mutation causes an increase in activity. On the other hand R312L shows decreased activity for both HA and ManA^10^.We present our paper in the following manner: The methods used to perform molecular docking on three sugars (HA_2_, HA_4_, ManA_6_) to Smlt1473 as well as its various mutants and all-atom MD Simulations, the various analysis techniques for MD simulations, as well as the various wet lab techniques for various mutagenesis experiments are discussed in Section 2. In Section 3, we present results on each of our sugar molecules and the various analysis performed.

## 2 Methods

### 2.1 Molecular Docking

The X-ray crystal structure of Smlt1473 in both the apo form as well as in the substrate-bound form was previously determined^10^. However, the structure of the substrates bound to Smlt1473 is not well resolved. To setup the initial structure, the sugar structures were modelled using CHARMM-GUI *Glycan Reader & Modeler*^12, 13^. The various sugars that we choose are (i) HA_2_-2 units of hyaluronic acid (GlcNAc-β-(1,4)-GlcA),(ii) HA_4_-4 units of hyaluronic acid and (iii) ManA_6_-6 units of mannopyranuronic acid (ManA) because Smlt1473 is the first CAenzyme to show pH dependent multiple substrate specificity for these sugars^7, 14^. The point mutations on various residues on the protein were carried out using PyMol. Then, we generated a conformational ensemble of the sugars using OpenBabel^15^. Ligand docking was subsequently performed using the Rosetta flexible docking protocol^16, 17^. In low resolution docking, the ligand is first translated as a rigid body around the binding pocket of the protein followed by rotation of the ligand and translation of the protein and the ligand towards each other. Then, high-resolution docking is performed by cycles of sampling of side chain rotameric conformers coupled with small translational motions of the ligand. These steps were repeated until convergence to generate 5000 complex models. These models were first sorted based on total energy scores and the top 20% of the models with the lowest total energy scores were further screened for the lowest total interfacial energy which was selected as our docked state. We repeated this procedure for all the ligands and the protein for both WT and its mutants.

### 2.2 All-Atom Molecular Dynamics Simulations

First, we chose a pH of 5 for and and a pH of 9 for ManA_6_ as this is the pH at which Smlt1473 shows most optimal activity for these ligands^7, 14^. Then, we ran the PROPKA3.1^18, 19^ server to determine the p of the various residues on the protein to determine their ionization states. To allow for variability, we also conducted the simulation on both unprotonated and deprotonated forms of the residue that has a p nearest to the chosen pH(+/-0.10 of the chosen pH) for simulation. This led to protonated and unprotonated forms of D144 for HA simulations and K130 for our ManA simulations. The protonated residues vary between the sugars mainly because of the varying sugar interactions at a different pH. We generated input scripts for MD simulation from CHARMM-GUI *Solution Builder*^20, 21^ and performed standard NTER and CTER patching for the terminal protein residues. Next, we assigned the ionization states of the various residues on the protein according to our PROPKA results and added water, potassium and chloride ions to neutralize the system. The periodic boundaries were setup resulting in a total of ∼40,000 atoms.

We ran the simulations in the GROMACS^22^ simulation package using a CHARMM36m^23–26^ force field and TIP3P^27^ water model. Minimization was performed using the Steepest Descent algorithm. A LINCS^28^ algorithm in the minimization, equilibration, and production constrained the hydrogen atoms. Equilibrations were run in the NVT ensemble and production runs in the NPT ensemble with a Nosé-Hoover thermostat^29^ at a temperature of 303.5 K. The NPT productions were run with a Parrinello-Rahman barostat fixing the pressure to be 1 bar with a compressibility of 4.5 X 10^−5^ bar^−1^ and a coupling constant of 5 ps. Position restraints were set on both the protein and the glycan atoms during the minimization and equilibration steps. During the production run, the position restraints were removed. We ran 5,000 steps of minimization and 125,000 steps of equilibration with a 1 fs time step. We further ran 375,000,000 steps of production runs with a 2-fs time step. This results in approximately 750 ns of production run for each ligand with 2 replicas comprising of both the protonated and unprotonated forms of a chosen residue.

### 2.3 Analysis of Simulations

Various analyses of the simulation trajectories were done in python. Root-mean-squared deviation (RMSD), fluctuation (RMSF), and fluctuations of the distance between specific residues were analyzed using MDAnalysis^30^ in python.

To determine the stable states from the simulation, we collected data from the simulation trajectories and performed clustering on the data using a simple K-Means algorithm (KMeans)^31^ and a deep-learning based algorithm Gaussian mixture variational autoencoders (GMVAE) ^32, 33^. To convert the raw simulation trajectory into a usable machine learning (ML) form, we collected the distances between the C_α_ carbons on the closest residues on the protein and the center of mass of each of the individual sugar units. The set of residues chosen here are based on the number of residues that are close to the center of mass of the sugar by a certain distance which we tuned to get a good enough value for our loss function. The set of these distances is used as input data for our ML algorithms. Thus, we show the applicability of entire protein-ligand simulation trajectory using two ML-based algorithms: KMeans and GMVAE.

Once we have these various clusters, binding energies of the clusters are obtained using the gmx-energy function of GROMACS. We then analyze the various interactions in each of these clusters. Hydrogen bonds are analyzed using the Visual Molecular Dynamics(VMD)^34^ Hbonds plugin. Salt-bridges and non-polar interactions are analyzed using MDanalysis using a distance-based cutoff between heavy atoms of the residues. The various snapshots from the simulations are obtained using PyMOL^35^.

### 2.4 Molecular Biology

An *E. coli* codon-optimized nucleotide sequence of Smlt1473 (GenBank accession number CAQ45011) was synthesized and cloned into pET-28a(+) using NcoI/XhoI restriction sites (GenScript). The coding sequence contains N- and C-terminal hexahistidine tags suitable for our protein purification method via affinity chromatography. The plasmid was transformed into *E. coli* DH5a for DNA isolation and *E. coli* BL21 for protein expression. Point mutations K56A, R107G, and R164G were made using the Q5 Site-Directed Mutagenesis Kit (NEB). Primers were designed using NEBaseChanger™. All sequences were verified by DNA sequencing (Eurofins).

### 2.5 Protein Expression and Purification

Mutant versions of Smlt1473, verified via DNA sequencing, were transformed into *E. coli* BL21 cells. Cells were grown in 10 mL of LB with kanamycin (50 μg/mL) overnight at 37 □ and shaking at 200 rpm. Cells were harvested by centrifugation at 3,000 *g* for 10 min. and then resuspended in 1 mL of fresh LB media. 1 mL of the cell suspension was used to inoculate a flask with 250 mL of LB. Flasks were incubated at 37 □ shaking until an OD_600_ between 0.6-0.8 was reached. Protein expression was induced using 1 mM isopropyl β-D-1-thiogalactopyranoside (IPTG) and then flasks incubated at 20 □ shaking for 16-18 h. Cells were harvested by centrifugation at 5,000 *g* for 15 min., supernatant was discarded, and cells were resuspended in lysis buffer (50 mM Tris-HCl, 100 mM NaCl, 5% v/v glycerol) prior to sonication. 30 mL of lysis buffer was used for 250 mL of expression culture. Sonication was performed with 20 s pulses for 20 min. After sonication, mixture was clarified by centrifugation at 10,000 *g* for 15 min. The supernatant containing protein was collected and insoluble fraction discarded. Supernatant containing protein was purified via immobilized-metal ion affinity chromatography (IMAC). The column was equilibrated with 5 column volumes of 10 mM imidazole/IMAC solution before adding the 0.2 μm filtered protein-containing supernatant. The protein was separated using imidazole elutions of 10, 30, 100, and 250 mM; 250 mM typically yielded the highest concentration of mutant Smlt1473. Each fraction was analyzed by SDS-PAGE with Precision Plus protein all-blue molecular weight standard (Bio-Rad) used as a molecular weight standard. The fractions with highest purity were then dialyzed in 2 L of 20 mM sodium phosphate buffer (pH 8) overnight in the 4 □ with stirring. Protein concentration was determined by measuring the absorbance at 280 nm and determining concentration using a calculated extinction coefficient of 74,495 M^−1^ cm^−1^. Protein aliquots were stored at −20 □ to be used in future experiments.

### 2.6 Enzyme Activity Assay

Polysaccharide lyases depolymerize polysaccharides via a β-elimination mechanism which results in double bond formation that can be detected by measuring absorbance at 235 nm^36^. For activity assays, the substrates used were polymannuronic acid sodium salt (BIOSYNTH) and high molecular weight (1.01-1.8 MDa) sodium hyaluronate (Lifecore Biomedical). Purified Smlt1473 (17.5 μg), both mutant and WT forms, were added to either ManA or HA (1 mg/mL) in a final reaction volume of 350 μL in UV-Star 96 well plates. ManA reactions were done at pH 9 using 30 mM Tris buffer and HA reactions were done at pH 5 using 30 mM sodium acetate buffer as pH for optimal enzyme activity against these substrates was previously determined^7^. Absorbance at 235 nm was measured at t = 0 and t = 10 min. Specific activity was calculated with one unit of enzyme activity being defined as an increase in absorbance at 235 nm of 1.0/min. Smlt1473 mutants H221F and Y222F were used as controls as their activity against these substrates has been previously reported^37^. Data were analyzed with respect to average WT values to determine the percent change in activity for each mutant against a given substrate.

## 3 Results

### 3.1 HA_2_ Binding to Smlt1473

First, we performed simulations for HA_2_ bound to Smlt1473 in water for both the WT and H221F mutation. All simulations show that the HA_2_ complex is bound to the enzyme during most of the simulations. Also, our simulations show that the RMSD of the protein backbone is similar between replicas and mutant (Figure S1). The RMSD of the protein backbone fluctuates around 2.5 □ which is a much lower value than the RMSD value of 5 □ shown by apo simulations^8^. The protein takes 100-200 ns to stabilize its structure. For the WT simulation (set 2) Figure S1B, there is an increase in the RMSD at the end of the simulation due to the gradual opening of the flexible loop and the adjacent helical region (red and blue region in Figure 1B). To highlight this effect, we have also plotted the RMSD of the loop region (Figure S1). The RMSD of the loop region shows a trend which is similar to the RMSD of the overall backbone which means that the changes in RMSD of the flexible loop region dictates the changes in the RMSD of the backbone. The RMSD of the sugars are shown in Figure S2.

**Figure 1:**
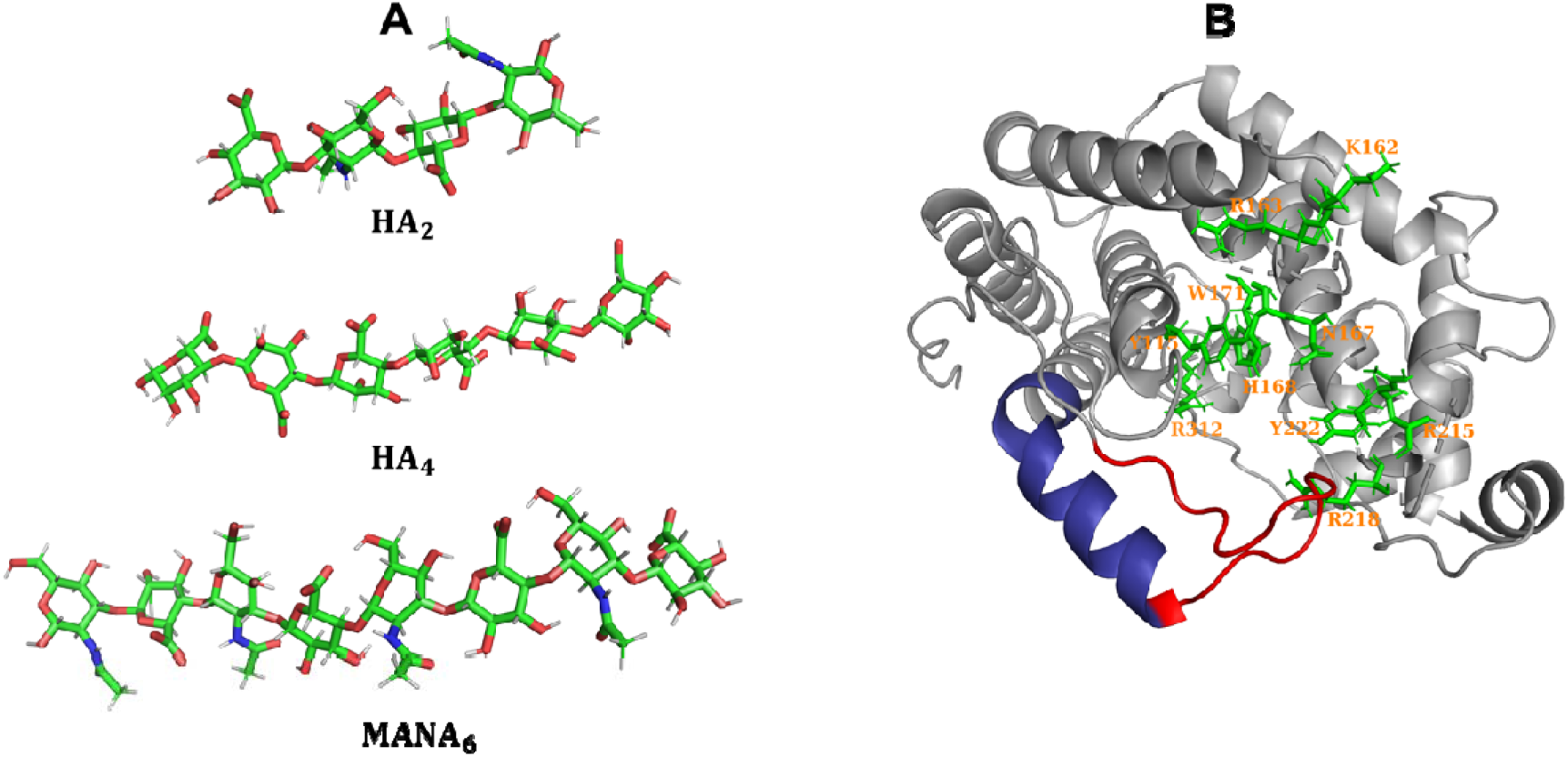
(A) Structures of the various sugar ligands discussed in this paper. (B) Structure of Smlt1473 with the entrance loop colored in red, the flexible helical region colored in blue and various residues important for sugar binding shown in green.

Next, we calculated the RMSF for all the protein residues. We have color coded Smlt1473 with the respective fluctuations for the various residues in Figure 2A for WT (set 1). The color coded Smlt1473 for WT (set 2), H221F (sets 1 and 2) as well as the fluctuation values for all sets are shown in Figure S3 and S4, respectively. The maximal fluctuations occur for the WT simulations (Figure S4), as the RMSF values for H221F (set 1) are much less than WT (set 1) and WT (set 2).

**Figure 2:**
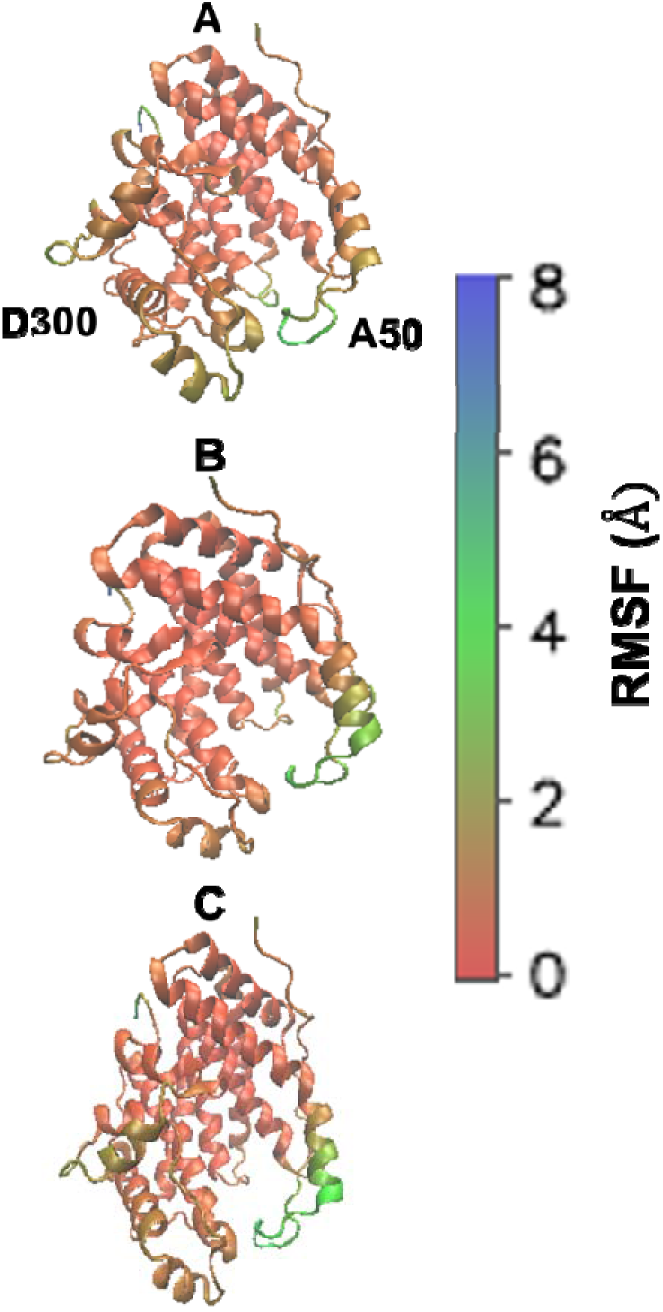
RMSF for Smlt1473 WT replicas for (A), (B), and (C).

As highlighted in Figures 2, S3, and S4, the regions of highest fluctuations are near the flexible loop region near the binding pocket (near residue 50) and towards the end of the protein distal to the binding pocket near residue 300. From Figure S5, we can see the pairwise fluctuation of distance between the catalytic residues are stable except for residue 42 which is located near the binding pocket (green region near the binding pocket in Figure 3).

**Figure 3:**
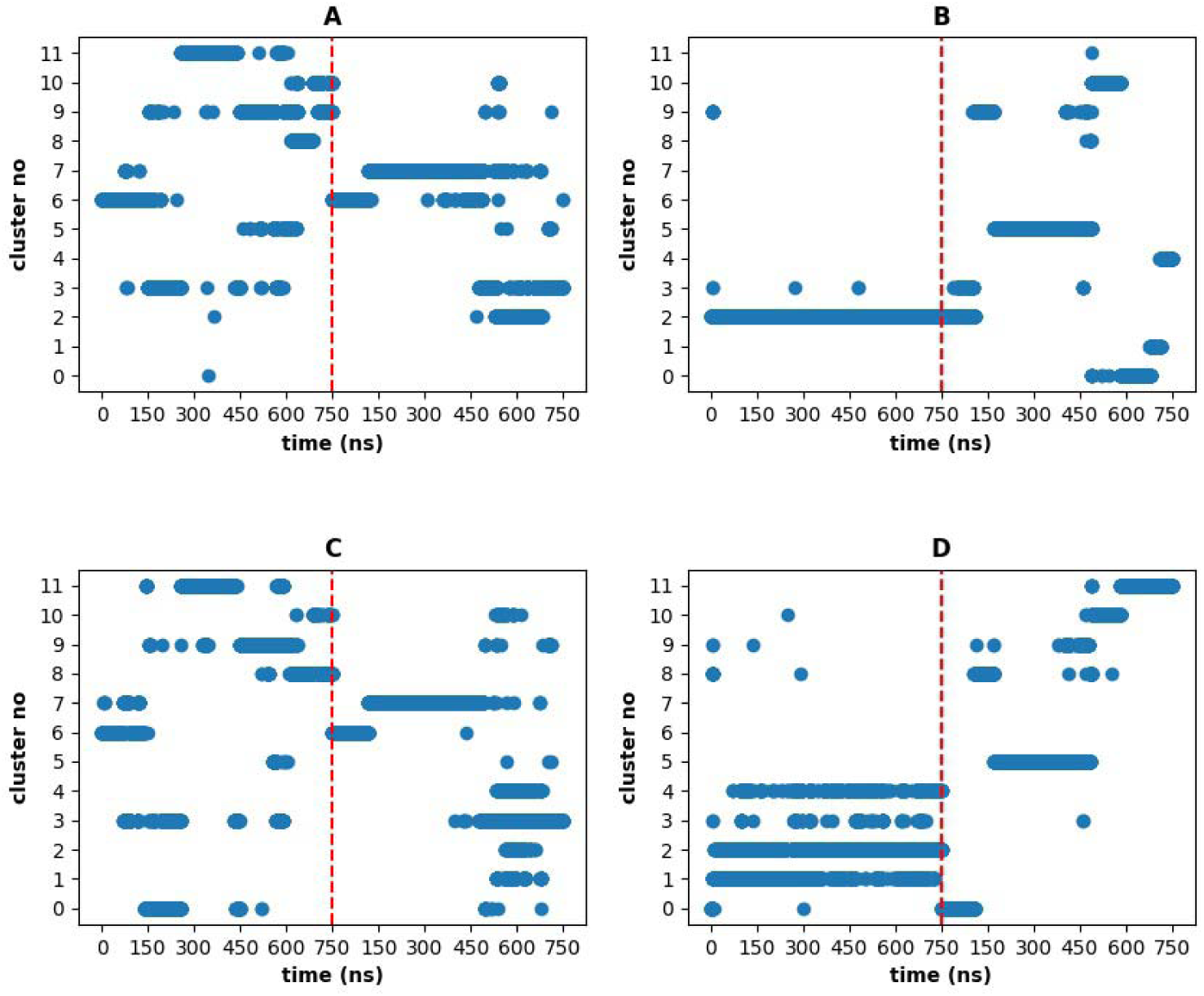
(A)-WT and (B)-H221F denote the various clusters predicted by the KMeans for the various structures of the protein-sugar complex. (C)-WT and (D)-H221F denote the various clusters predicted by GMVAE for the various structures. The red line in between separates the two sets for the simulations with HA_2_.

Next to classify the diverse conformational states of HA_2_ bound to Smlt1473, KMeans and GMVAE (Fig. 3) are used to determine the clusters focusing on the center of mass of the sugar and its closest distances to important protein residues. In a majority of the cases, GMVAE gives improved and insightful cluster information as compared to KMeans due to the overreliance of the KMeans to classify data based on a Euclidean distance metric as compared to clustering in a low dimensional latent space by GMVAE. The WT clustering results are mostly similar. For H221F simulation, GMVAE is able to find fluctuations between the various states for the first set of simulations. This fact is highlighted in the distance maps in Figure S6C as well which shows fluctuations in colors at residues 163, 167, 168, 215, 218, 222, and 312. From these various clusters, we can also find out the most dominant interactions in the most dominant clusters. We have also individually determined the major cluster energies in Table S1 and presented bar graphs in Figure S7.

To distinguish the various clusters, we determined the interaction energies of the most important residues in the major clusters as shown in Figure 4 and Tables S2 and S3. Figure 5 displays the most important interactions in the major clusters. The interaction energy values are well separated in each of the clusters showing the efficiency of our clustering algorithm in identifying clear clusters. The results for the other clusters are given in Tables S4 and S5 and Figures S8 and S9.

**Figure 4:**
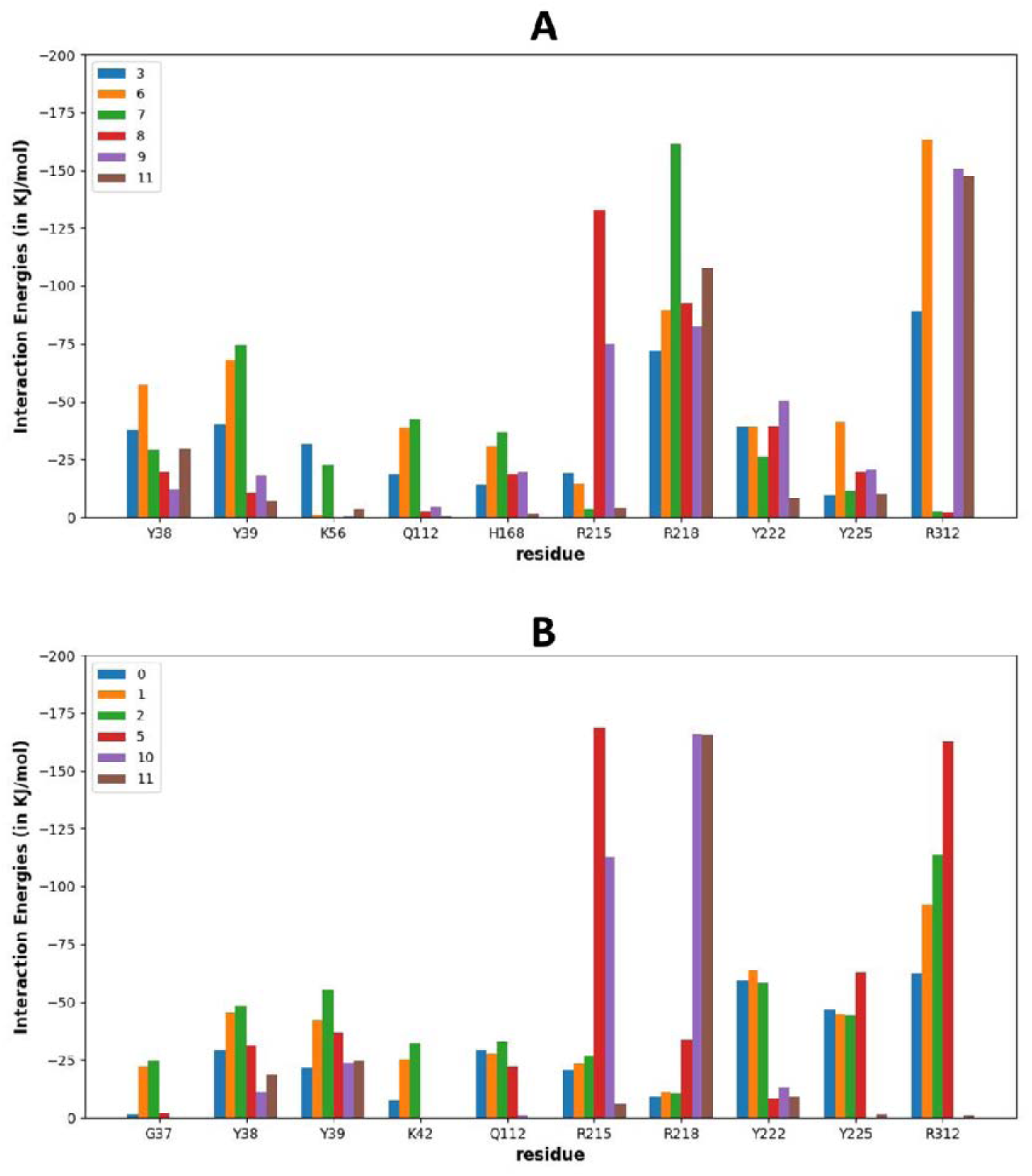
The average interaction energies of each of the highest interacting residues to the sugar from (A) WT simulation and (B) H221F mutant for the highest populated clusters with HA_2_. The detailed interaction energy values are given in Tables S2 and S3.

**Figure 5:**
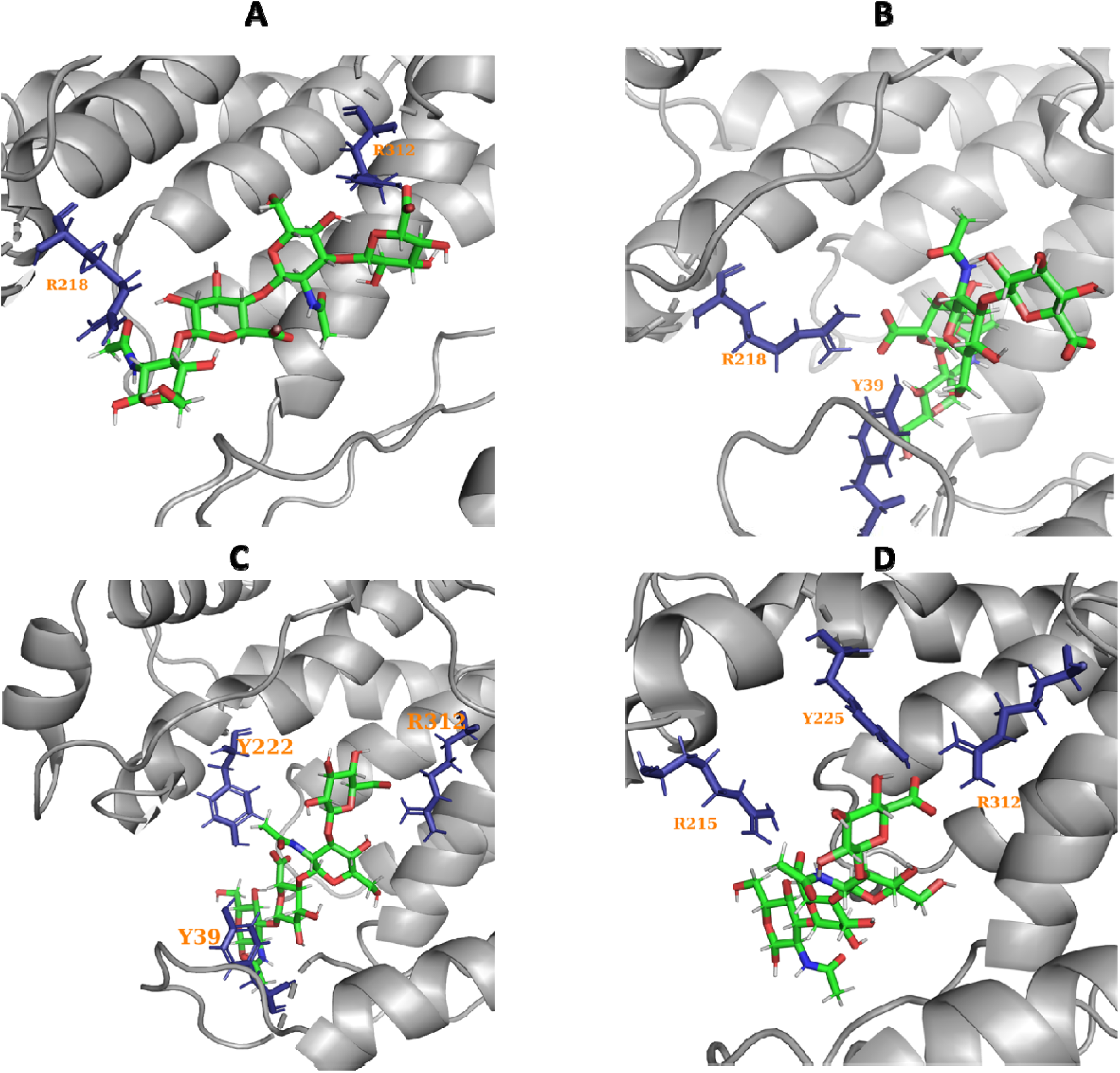
The two highest populated clusters with the respective highly interacting residues to the sugar from WT simulation ((A) and (B)), H221F mutant ((C) and (D)). For WT simulation, it is clusters 3(A) and 7(B) and for H221F simulation it is clusters 2(C) and 5(D).

In Figures 4A, 5A, and 5B, the most important interactions between the WT protein and the sugar for the two majorly populated clusters are shown. R218 forms strong salt bridges in both of these clusters, while R312 only forms stable salt bridges in cluster 3. Additionally, Y39 only forms strong hydrogen bonds in cluster 7. Other important residues that form non-polar contacts are G37, N109, N110, D111, F114, P161, K162, R163, N167, A214, Q217, L220, H221, and D224.

In Figures 4B, 5C, and 5D, the most important interactions between the H221F mutant and the sugar for the majorly populated two clusters. R312 forms stable salt bridges for these two clusters and R215 forms salt bridges for cluster 5. Additionally, Y39 and Y222 form stable salt bridges for cluster 2, while Y225 forms stable salt bridges for cluster 5. The major non-polar contacts in the H221F type simulation are R34, A35, I36, G37, Y38, Y39, T40, D41, K42, A43, V46, Q54, N110, D111, F114, Q118, N167, A214, Q217, L220, F221, D224, and P311.

In addition, we have further investigated the changes due to the H221F mutation. These major changes are shown in Table 1 and Figure 6. Due to the mutation, there is an increase in the overall interaction energy of the enzyme to the ligand. The average binding energy of the WT and H221F is −505±1 KJ/mol and −520±1 KJ/mol, respectively. In the H221F mutation, the R218 residue located near residue 221, which interacts weakly with the sugar compared to the WT. There are certain residues such as S310 and H168 that interact slightly weaker with the sugar as well due to the mutation. This causes a much stronger interaction of the sugar with other residues, i.e., G37, K42, R164, R215, Y222, Y225, and R312 in the mutant. In total, these changes result in more attraction of HA_2_ to the protein with the H221F mutant.

**Figure 6:**
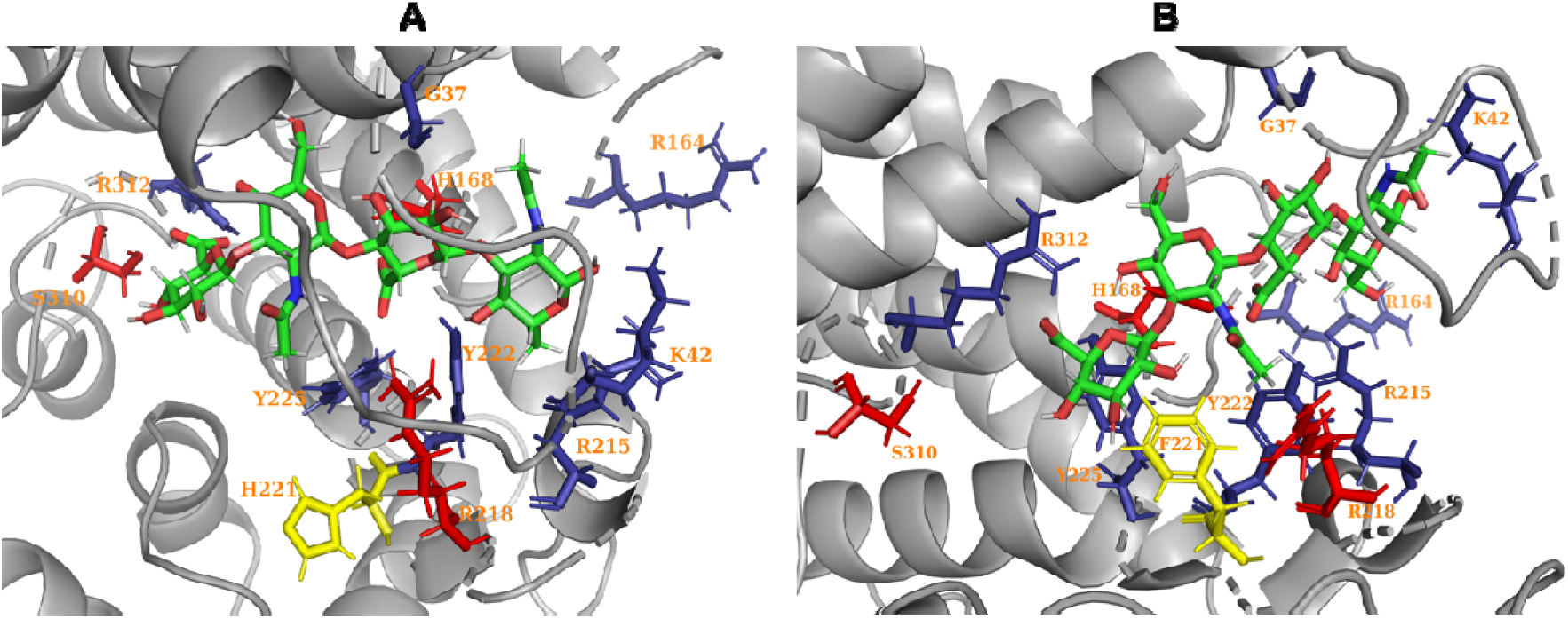
(A) WT structure and (B) H221F mutant with HA_2_. This figure highlights the key residues shown in Table 1 which shows major changes in the interaction energies upon mutation and leads to an overall decrease in interaction energy on the H221F mutation. The H221 and F221 are highlighted in yellow. The residues whose interaction energies decrease upon the mutation are highlighted in red and the residues whose interaction energies increase are highlighted in blue.

**Table 1:** The major changes in the sugar-protein interaction energies for the H221F mutation with HA_2_.

| Residue | WT(in KJ/mol) | H221F(in KJ/mol) |
| --- | --- | --- |
| G37 | -2.29 | -12.2 |
| K42 | -6.42 | -15.1 |
| R164 | -3.01 | -9.26 |
| R168 | -21.4 | -7.03 |
| R215 | -29.3 | -63.9 |
| R218 | -106 | -43.6 |
| Y222 | -32.6 | -40.5 |
| Y225 | -18.5 | -36.8 |
| S310 | -9.06 | -1.04 |
| R312 | -81.1 | -88.2 |

### 3.2. HA_4_ Binding to Smlt1473

To investigate the effect of a longer polysaccharide, we performed simulations for HA_4_ bound to Smlt1473 in water for both WT and H221F. Our simulations show that HA_4_ is bound to the enzyme during most of the simulations. The RMSD of the protein backbone in all of the simulations is similar to simulations with HA_2_ with the protein backbone stabilizing to around 2.5 □ as shown in Figure S10 which is a much lower value than the previous results shown by apo simulations^8^. The overall protein structure takes 100-200 ns to equilibrate with the WT and H221F, making them similar in protein stability. As we had shown for HA**_2_**, the changes in the RMSD of the sugar are dictated by the changes in the RMSD of the flexible loop and the adjacent helical region (residues 36-63). For example, the WT (set 2) simulation shows some structural changes related to the opening of the flexible loop region (near 250 ns). At around 450 ns, the blue helical region (Figure 1B) near the red flexible loop gradually starts to move out of the binding pocket and this causes the flexible loop region to open up towards the end of the simulation. The RMSD of the sugars are given in Figure S11.

We have also plotted the RMSF of the various amino acid residues in Figure S12. The fluctuations are the most dominant near the loop and the flexible helical region-1 (near residue 50, blue regions near active site in Figure S12). These WT simulations show more fluctuations near loop 1 as compared to the mutated simulations. The other region that shows elevated fluctuations is near the C-terminus (residue 300, see the green region not near the binding site in Figure S12).

As noted in previous experimental work^7, 8, 11, 14, 37, 38^, the loop region (specifically K42) is important for the enzyme activity and this region shows high fluctuation (Figures 2B, S12 and S13). We have also plotted the fluctuations of the distances between the various catalytic residues of the protein in Figure S14 to focus on the residues of dynamic importance. The distance between residue 42 and the catalytic residues show maximal fluctuations.

As with HA_2_, the sugar explores different bound conformational states within the binding pocket of the protein. To characterize the different bound conformational states of the sugar within the binding pocket of the protein, we performed different clustering algorithms and the various clustering results are shown in Figure S15 and S16. The major KMeans and GMVAE cluster energies are in Table S6 as well as the fraction of the percentage of each of the clusters. We have also represented the cluster binding energies as bar graphs in Figure S17. Our focus for the remaining of the manuscript will be the clustering from GMVAE as it provides more insightful clusters, but the SI also contains KMeans results for reference.

To distinguish the various clusters, the interaction energies of the top 10 interacting residues were calculated for the WT and the H221F mutant (Table S7 and Table S8 respectively). There is a clear distinction in the interaction energies of the various important residues with the sugar for each of the clusters. The values of these energies show clear distinction which shows that we have been able to generate well defined clusters using our GMVAE algorithm. We also show the cluster definitions in terms of the interaction energies as bar plots in Figure 7.

**Figure 7:**
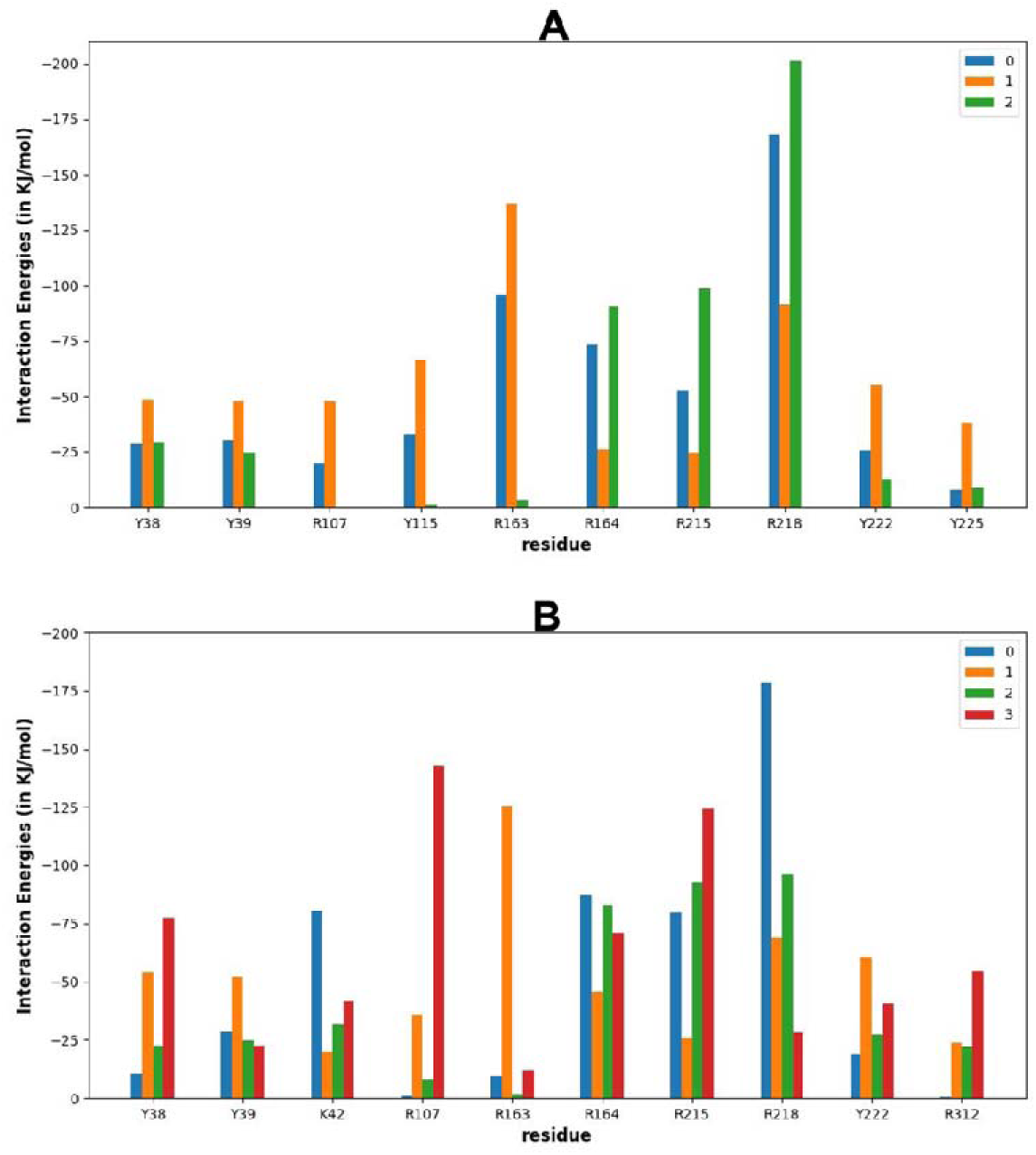
The interaction energies of the most important residues in the various stable clusters from the (A)WT simulation and (B) H221F simulation sets with HA_4_. The cluster numbers are color coded according to the legend. The detailed data about the various clusters are shown in Tables S7 and S8.

As shown in Figures 8A, 8B, and S18A, for the WT simulation, R107 forms stable salt bridges in cluster 1. R163 forms stable salt bridges in clusters 0 and 1. R164 and R215 form stable salt bridges in clusters 0 and 2. R218 forms stable salt bridges in clusters 0, 1 and 2. Y38, Y39, Y115, Y222, and Y225 forms hydrogen bonds with cluster 1. Other important contacts are G37, Y38, N109, N110, D111, F114, P161, K162, N167, A214, Q217, L220, H221, D224, and Q277.

**Figure 8:**
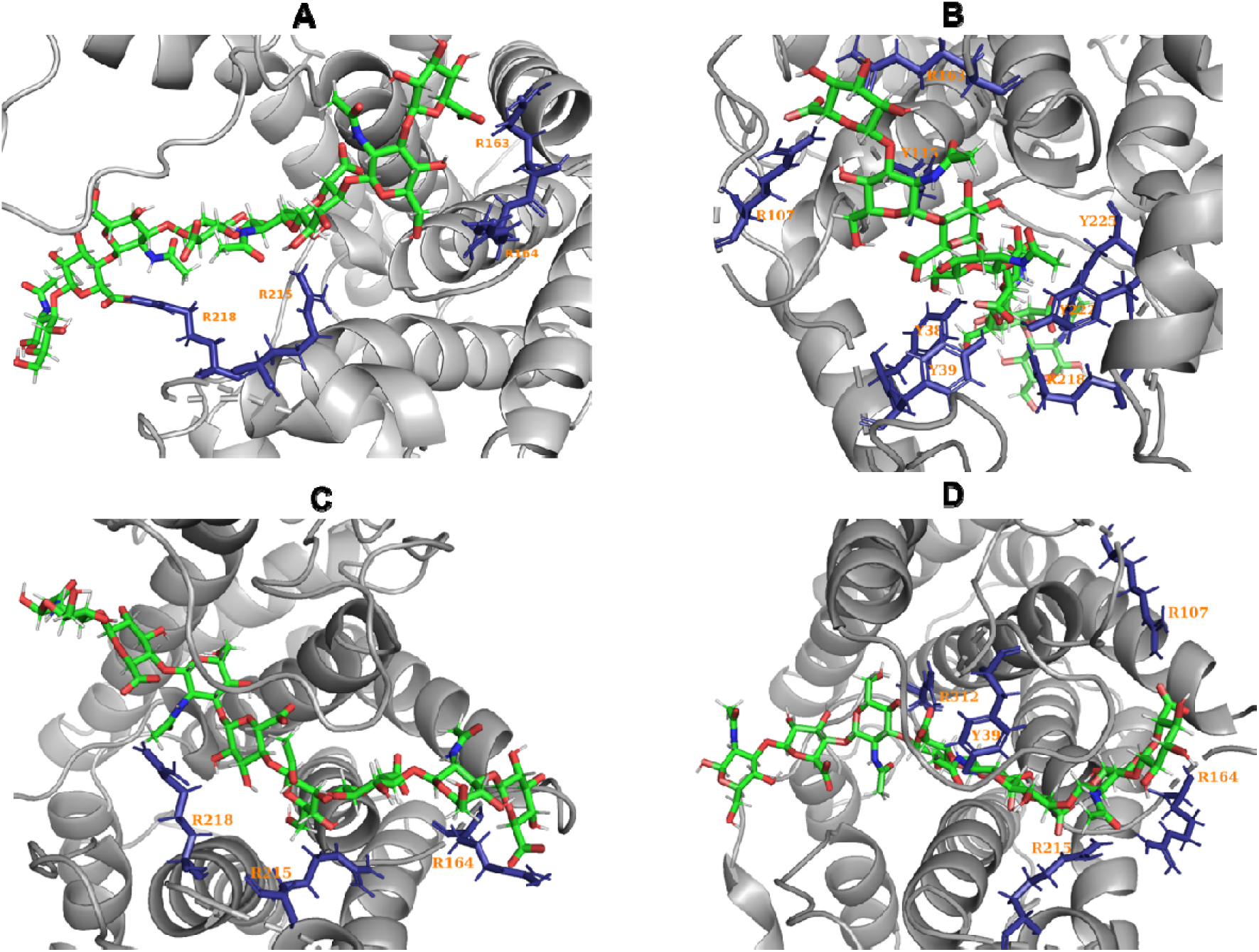
The highest 2 populated clusters with the respective highly interacting residues to the sugar from WT simulation ((A) and (B)) H221F mutant ((C) and (D)). For WT simulation, it is clusters 0(A) and 1(B) and for H221F simulation it is clusters 2(C) and 3(D).

From Figures 8C, 8D, S18B, and S18C, for the H221F mutant, K42 forms stable salt bridges in cluster 0. R107 and R312 form strong salt bridges in cluster 3. R163 forms a stable salt bridge in cluster 1. R164 and R215 form stable salt bridges in clusters 0, 2, and 3. R218 forms stable salt bridges in clusters 0, 1, and 2. Y38 and Y222 form stable salt bridges in clusters 1 and 3. Apart from this, we find that the H221F mutant also forms important non-polar contacts with other residues like and G37, Y38, D41, A43, G44, V46, L51, I106, V108, N109, N110, D111, F114, Q118, N166, N167, Y170, L211, A214, Q217, L220, F221, F224, Y225, A282, F308, H309, and P311 (set 2).

Overall, we found that the average total interaction energy of the WT is slightly lower (−747±2 KJ/mol) than the H221F (−750±2 KJ/mol). Although similar, on closely investigating the per residue interaction energy values, we see that the mutant is causing some long-range effects in the overall bound state. By looking at Table 2 and Figure 9, H221F is drastically reducing the interaction energy of R218 with the sugar which is located very close to 221 H/F(in yellow), as well as Y115 and R163, which are located a bit far away. This causes several residues that were interacting less with the sugar to interact more strongly with the sugar (Y38, K42, R107, R164, and R215). However, this causes a slight decrease in the overall interaction energy.

**Figure 9:**
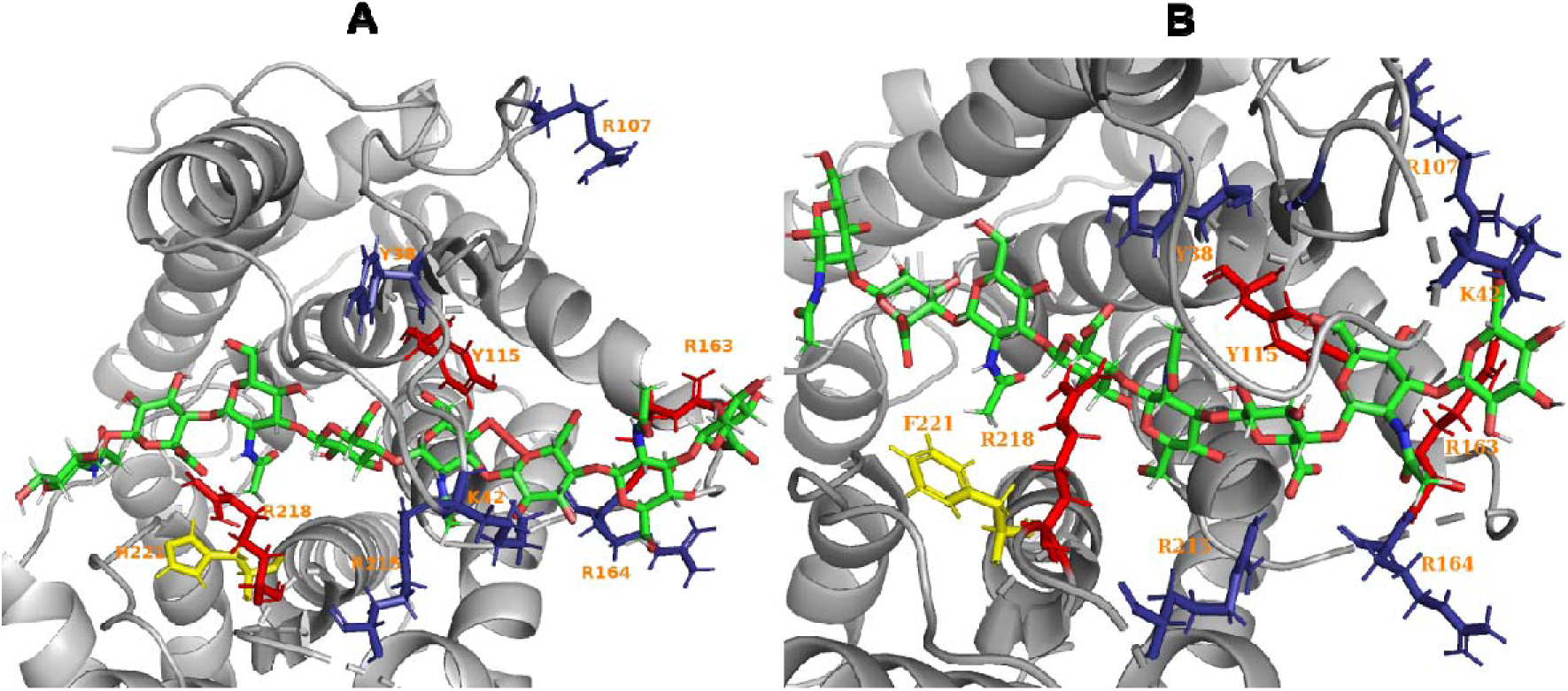
The major effects of the H221F mutation for simulations with HA_4_. The residues that interact more strongly due to the mutation are shown in blue and the residues that interact weaker are shown in red. (A) denotes the WT sample and B denotes the H221F mutant.

**Table 2:** The major changes in the interaction energies with HA_4_ due to Smlt1473 H221F mutation.

| Residue | WT(KJ/mol) | H221F(KJ/mol) |
| --- | --- | --- |
| Y38 | -36.7 | -50.4 |
| K42 | -16.3 | -42.7 |
| R107 | -25.9 | -72.9 |
| Y115 | -37.7 | -15.2 |
| R163 | -87.5 | -26.7 |
| R164 | -58.1 | -72.1 |
| R215 | -53.2 | -94.4 |
| R218 | -144 | -74.6 |

### 3.3. ManA_6_ Binding to Smlt1473

To complete our sugar binding study with Smlt1473, we performed simulations for ManA_6_ bound to the enzyme in water for the WT, H221F, and R312L versions of the protein. Nearly all simulations show that the ManA_6_ remains bound to the enzyme. As with the other sugars, all simulations show an overall backbone RMSD of the protein that is similar (Figure S19). The protein backbone takes around 150-200 ns to stabilize its backbone RMSD and fluctuates around 2-3 Å which is a much lower than the previous results shown by apo simulations^8^. As discussed for the previous two sugars, the structural changes in the overall backbone of the protein are mainly dictated by the structural changes in the loop 1 residues 36-63 as shown in Figure S20.

In Figure 2C, S21, and S22 the RMSF of the protein and its effect under various mutations is shown for each residue. The maximal RMSF for the WT simulations is in the region of loop 1. The next most elevated RMSF is the R312L mutation which shows a higher RMSF value near loop 1 as compared to the H221F mutation. This fact is further highlighted in Figures 2C and S22. We see that the proteins in Figure S21A (WT set 2) and S21E (R312L set 2) show the maximal RMSF near the binding pocket region. All systems show an elevated fluctuation near the C-terminus region of the protein around D300. The fluctuations of the distances between the various catalytic residues show residues located near the residue 42 shows the maximal fluctuations (Figure S23) like for HA_2_ and HA_4_.

We also performed clustering for the simulation trajectories and presented the results and comparisons in Figures S24. The comparisons of the clusters with the distance plots are shown in Figure S25. The binding energies for the various clusters by our clustering algorithms are shown in Figure S26 with detailed values in Table S9 as well as the fraction of residence of each cluster in Table S10. Again, for quantitatively distinguishing the various clusters, we show the major interactions energies occurring for the 10 highest interacting residues. A bar graph is shown in Figure 10 for each cluster in terms of interaction energies with the most important residues for the WT, H221F, and the R312L mutant. The interaction energies are well separated for these clusters highlighting the efficiency of our clustering algorithm. The detailed interaction energies for each set of residues are shown in the SI (Tables S11-S13).

**Figure 10:**
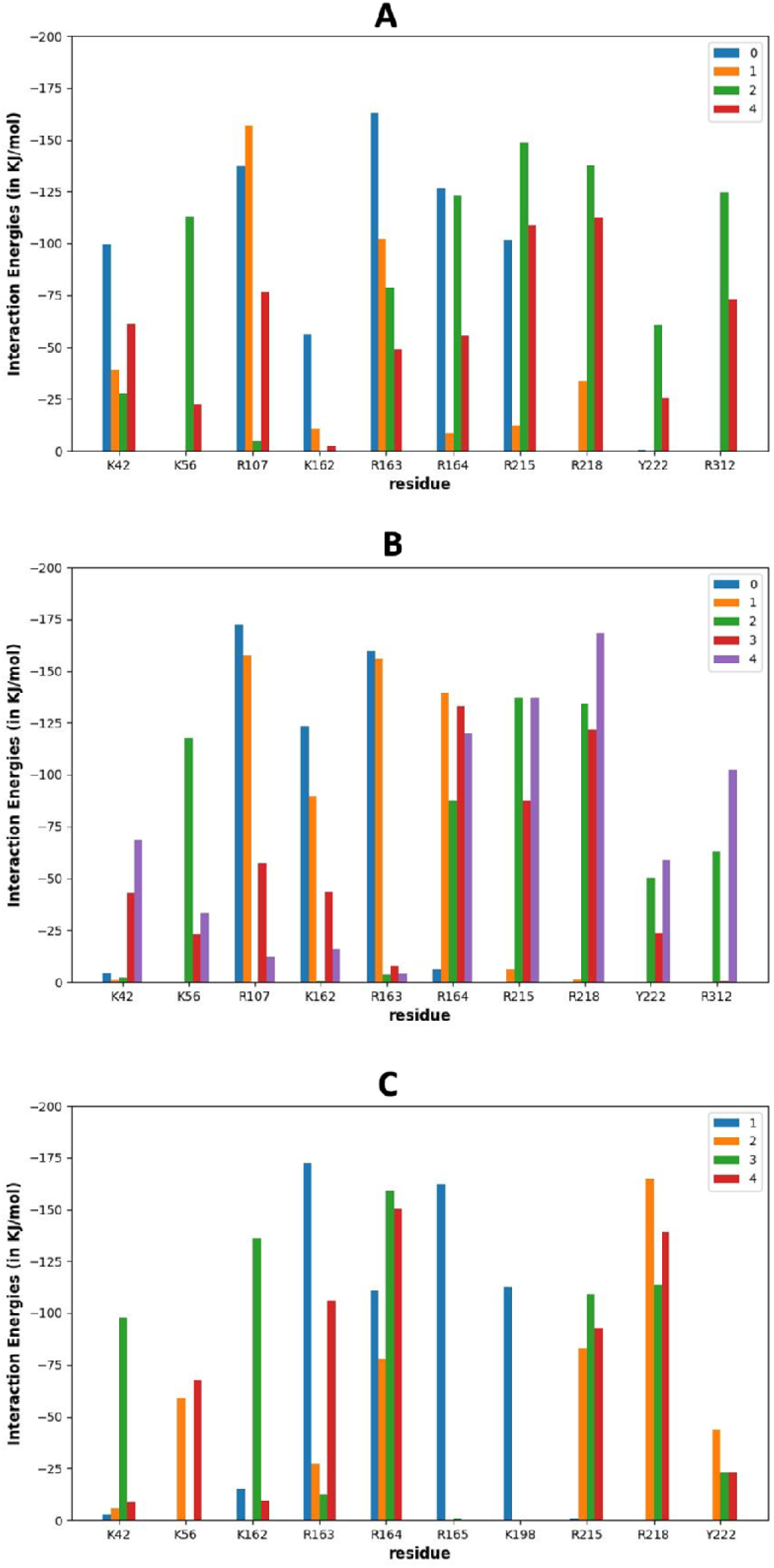
The various interaction energies by the various residues for the various clusters by the various residues. (A) shows the data for the WT simulation, (B) shows the data for the H221F mutant, and (C) shows data for the R312L mutant. The detailed data is shown in tables S11, S12, and S13.

Snapshots for ManA_6_ bound to WT and two mutants are shown in Figure 11A, 11B, S27A and S27B with important residues highlighted. K42 salt bridges stabilize clusters 0 and 4. Salt bridges at K56 and hydrogen bonds at Y222 stabilize cluster 2. R107 forms salt bridges in clusters 0, 1, and 4. K162 forms salt bridges in cluster 0. R163 forms stable salt bridges at clusters 0, 1, and 2. R164 and R215 form stable salt bridges at clusters 0, 2, and 4. R218 and R312 form stable salt bridges at clusters 2 and 4. Non-polar contacts that N55, N110, D111, F114, Q118, P161, R165, N166, N167, P210, L211, A214, Q217, H221, D224, and P311.

**Figure 11:**
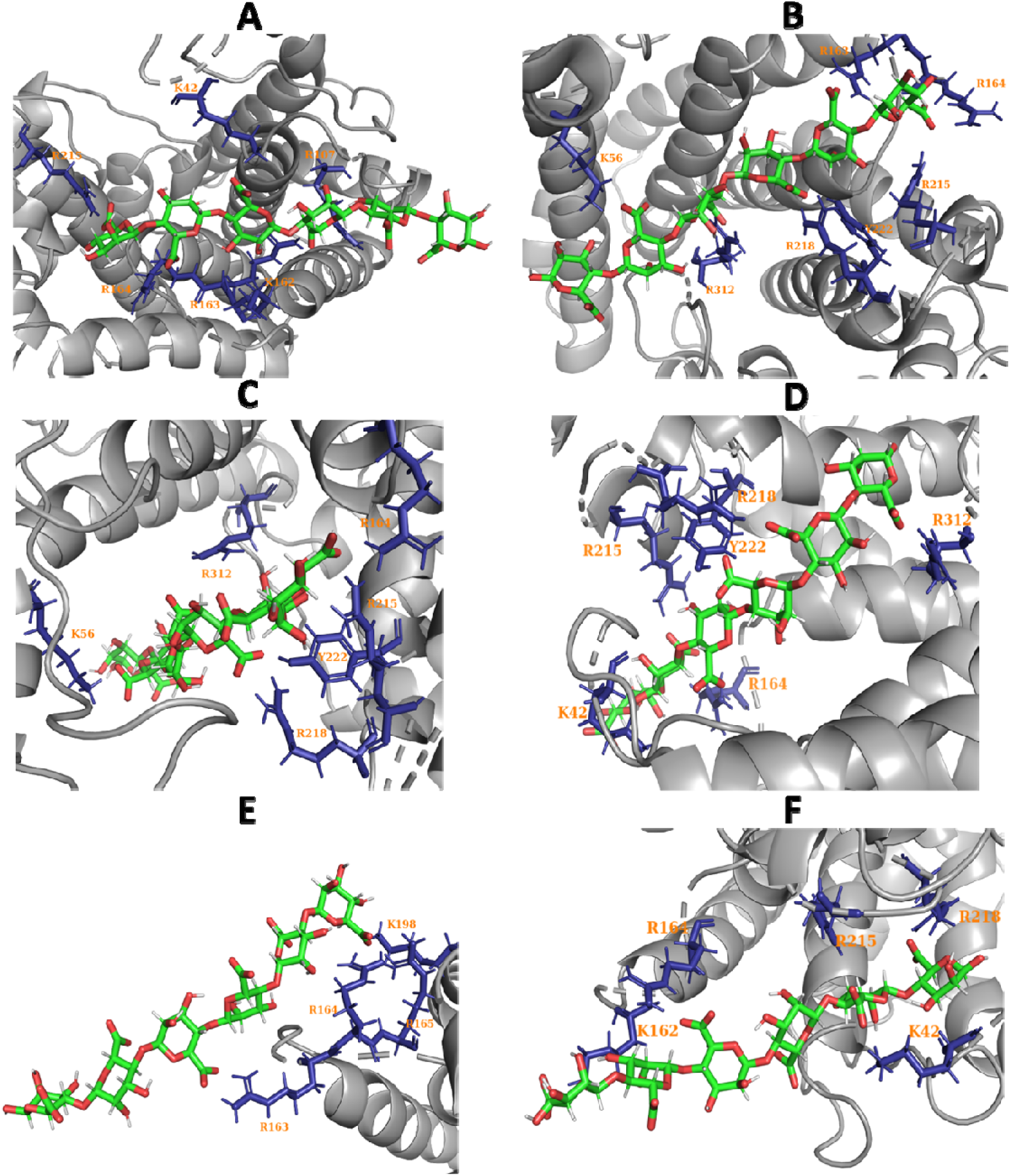
The two highest populated clusters with the respective highly interacting residues to the sugar from WT simulation ((A) and (B)), H221F mutant (C) and (D)), and R312L mutant (E) and (F). Here are the cluster codes: (A)- WT cluster 0, (B)- WT cluster 2, (C)- H221F cluster 2, (D)-H221F cluster 4, (E)- R312L cluster 1, (F)- R312L cluster 3.

From Figures 11C, 11D, S27C, and S27D, we can analyze the interactions stabilizing ManA_6_ to the H221F mutant. K42 stabilizes cluster 4 with salt bridges. K56 forms stable salt bridges in cluster 2. R107 forms stable salt bridges in clusters 0, 1, and 3. K162 and R163 form stable salt bridges at clusters 0 and 1. R164 forms stable interactions at clusters 1, 2, 3, and 4. R215 and R218 form stable salt bridges at clusters 2,3, and 4. R312 forms stable salt bridges and Y222 forms hydrogen bonds at clusters 2 and 4.

Figures 11E, 11F, S27E, S27F, and S27G provide information on the interactions stabilizing ManA_6_ to the R312L mutant. K42 and K162 form stable salt bridges in cluster 3. K56 forms stable salt bridges at clusters 2 and 4. R163 forms stable salt bridges in clusters 1 and 4. R164 forms stable salt bridges at clusters 1, 2, 3, and 4. R165 and K198 form stable salt bridges in clusters 1. R215 and R218 form salt bridges at clusters 2, 3, and 4.

To further investigate the changes brought by the H221F and R312L mutations, Table 3 and Figure 12 show the major changes in the ManA_6_ interaction energy with the enzyme. From these results, ManA shows differences due to mutations compared to HA. There is a major decrease in the overall interaction energy for H221F (−779 ± 2 KJ/mol) as compared to the WT (−840 ± 2 KJ/mol) simulations as opposed to the increase which we observed in the case of the H221F simulations for the HA sugar. The major cause can be attributed to R218 which was previously interacting weakly in the presence of the F221 residue for the HA sugar and for the H221F mutant now interacts very strongly in presence of ManA sugar. Apart from this, K162 tends to interact strongly as well. This causes several other residues (K42, K56, R107, R163, R164, R215, R218, and R312) to interact less strongly with ManA. This opposing effect due to the ManA sugar might be one of the main reasons why there is increase in activity due to the H221F mutant due to the ManA sugar as compared to the HA sugar unit^11^.

**Figure 12:**
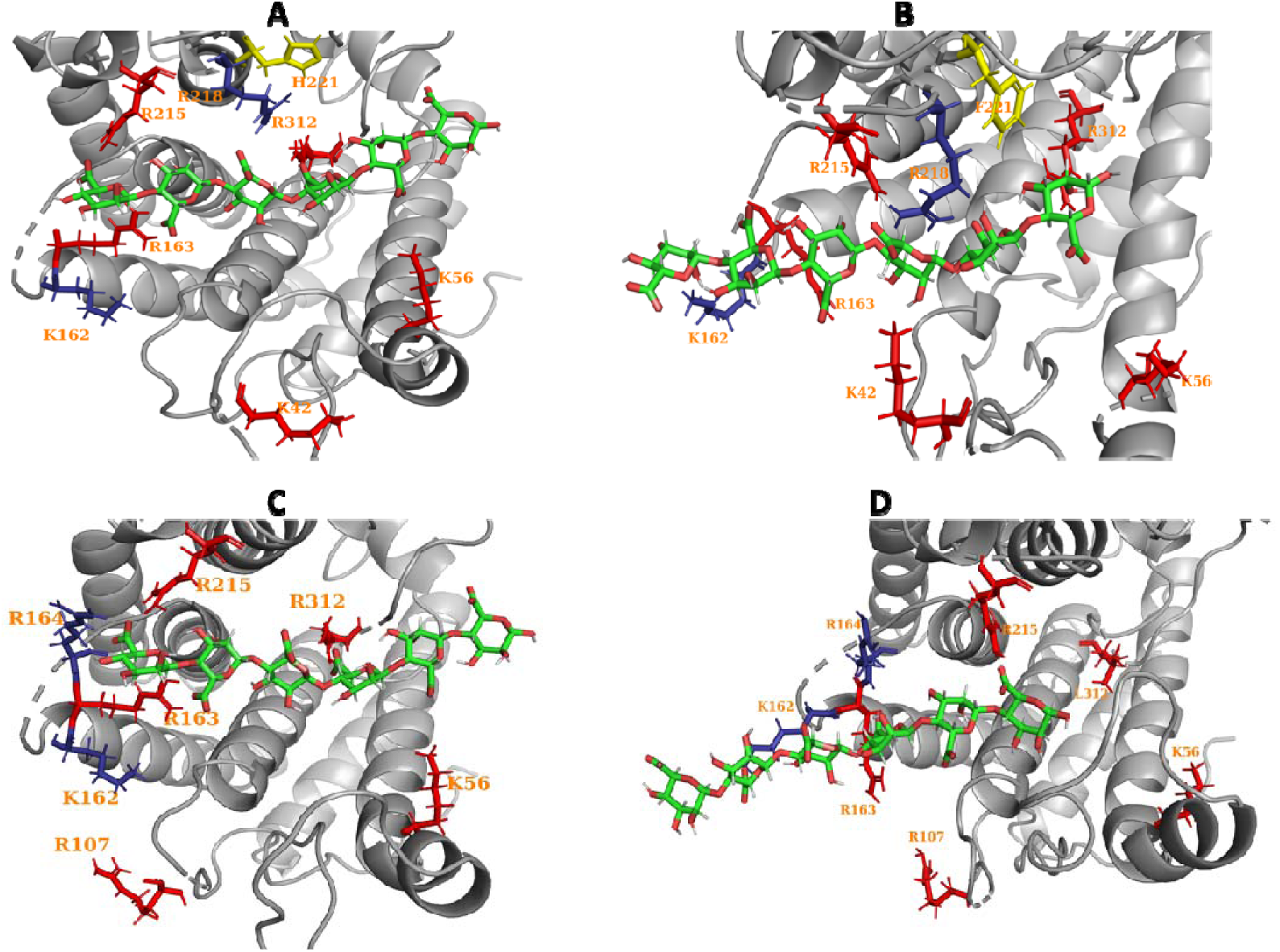
The residues which show a major change in interaction energy with ManA_6_. A and B compare the WT vs the H221F mutant respectively. C and D show the WT vs the R312L mutant respectively. The residues colored in red are those that show decrease in interaction energy upon the mutant. The residues shown in blue are those that show an increase in the interaction energy.

**Table 3:** The major residues which show a significant change in interaction energy of ManA6 with either the H221F or R312L mutation.

| Residue | WT (KJ/mol) | H221F (KJ/mol) | R312L (KJ/mol) |
| --- | --- | --- | --- |
| K42 | -55.6 | -35.4 | -46.4 |
| K56 | -67.5 | -40.1 | -15.9 |
| R107 | -56.7 | -54.3 | -8.70 |
| K162 | -20.8 | -40.1 | -66.5 |
| R163 | -108 | -43.7 | -76.5 |
| R164 | -121 | -103 | -134 |
| R215 | -129 | -96.8 | -70.9 |
| R218 | -85.8 | -113 | -87.8 |
| R312 | -76.3 | -55.9 | -0.138 |

The effects on ManA_6_ binding are easy to see due to the R312L mutation. The overall binding energy due to R312L is −666 ± 1 KJ/mol. The R312 mutation disrupts the stable salt bridge interaction between R312 and the sugar destabilizing the ManA sugar which causes a sharp decrease in the overall interaction energy of the enzyme to the sugar. This causes a sharp decrease in the interaction energies of many of the residues like K42, K56, R107, R163, R215, and R312 as well. Although this mutation causes an increase in the interaction energies of K162, R164, and R218, the overall effect is still to decrease the overall interaction energies of R312L mutation.

### 3.4 Effects of mutations to K56, R107, and R164 on Smlt1473 activity

Based on computational simulations, various residues with the highest interaction energies depicted (Figures 4, 7, and 10) were found to be important for ManA and HA binding. Many of these residues have already been explored in previous works, however K56, R107, and R164 were newly predicted by simulations done in this study to be important for sugar binding^7, 11, 14, 38^. To explore these new residues of interest, point mutations were made to Smlt1473 and activity of each mutant was experimentally determined against ManA and HA. Where the residue is located in the crystal structure of Smlt1473 (helix or loop), as well as maintaining the secondary structure of the protein were taken into consideration when choosing the amino acid to mutate to at these residues. R164 and R107 are both located within loop regions, so in both cases, arginine (positive and polar) was mutated to glycine (neutral and nonpolar). K56 is located within a helix and therefore, lysine (positive and polar) was mutated to alanine (neutral and nonpolar). K56A, R107G, and R164G were all tested for activity against both ManA and HA using absorbance at 235 nm. Figure 13A shows that K56A and R164G mutations did not lead to changes in activity against ManA w.r.t WT, however, the R107G mutation led to a significant decrease in activity against ManA. Additionally, Figure 13B shows that opposing the results for ManA, the K56A mutation led to a significant decrease in activity against HA w.r.t. WT. Similar to results for ManA, R164G did not result in any change in activity against HA. Interestingly, R107G showed a significant decrease in enzyme activity against HA as well, however, to a much greater extent than for ManA. H221F and Y222F were used as controls for the assay as their activities against ManA and HA had been previously determined^7, 14^. These data show that computational simulations modeling interactions between a protein and its substrate can provide valuable insight for determining residues that are most important to general binding, but also for substrate specificity. The models are valuable in that they can predict residues to explore without the intense burden of experimentally generating thousands of mutants at random.

**Figure 13:**
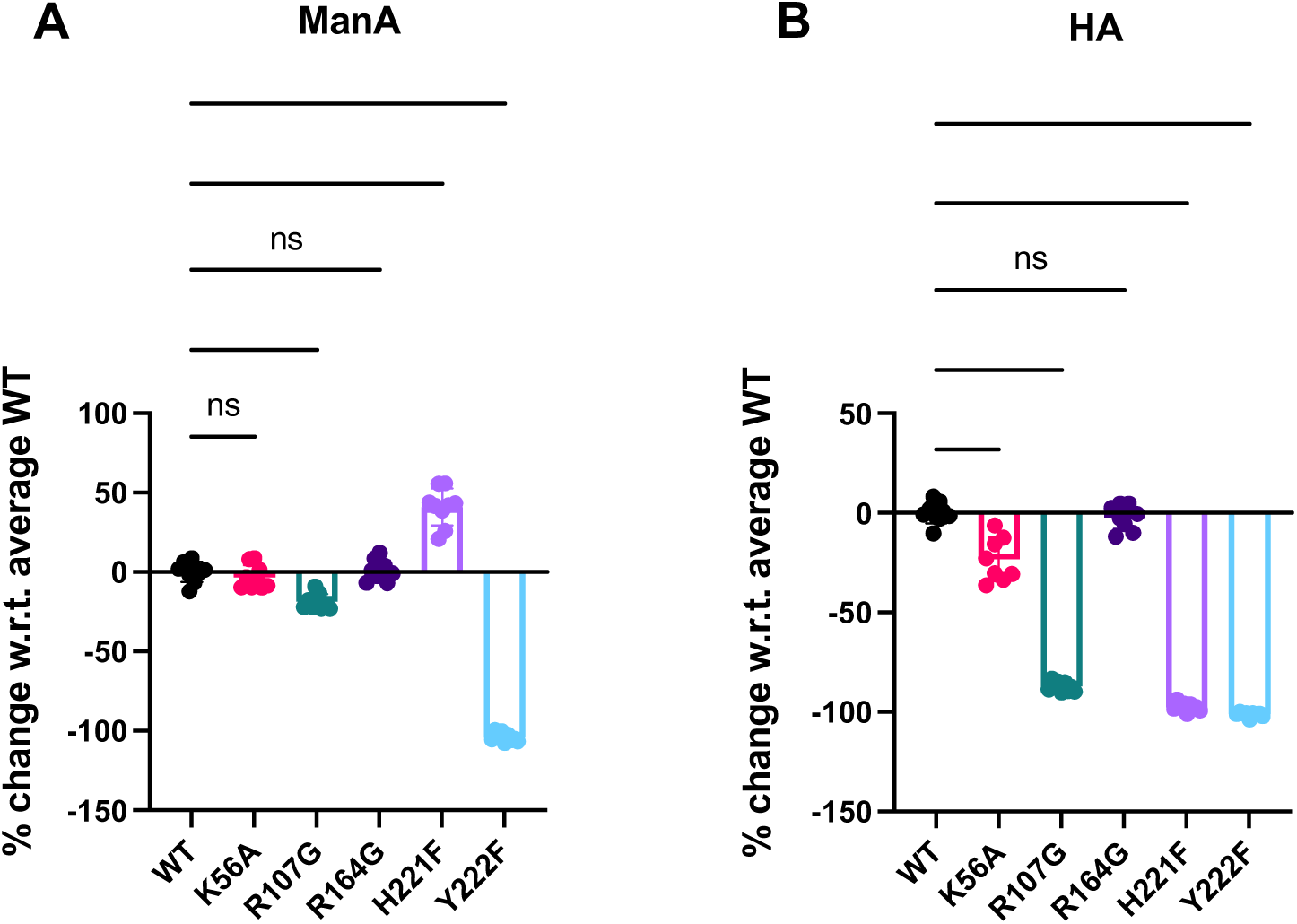
Purified WT and mutant Smlt1473 (17.5 μg) was added to 1 mg/mL HA in 30 mM sodium acetate at pH 5 or ManA in 30 mM Tris at pH 9 in a total reaction volume of 350 μL. Absorbance at 235 nm was monitored to determine specific activity for WT and mutant Smlt1473 against ManA and HA. Percent change in activity with respect to WT was calculated for each mutant against (A) ManA and (B) HA.

## Discussions/Conclusions

We have run MD simulations on Smlt1473 and its various mutants for a variety of sugars: HA_4_, HA_2_, and ManA_6_ by docking these sugars to the protein binding pocket and have studied the microscopic details of the interactions as well the dynamics of these sugars. In previous works, we have looked at the effect of mutations as well as pH on the activity for this enzyme^8, 11^. In our study, we have connected thermodynamics of the sugar binding to the previously determined activity of this enzyme by starting from its crystal structure. We have also developed a method to cluster protein-ligand simulation trajectories into different states. The sugars are mostly stable inside the binding pocket during the 750 ns simulation timescale. The protein stabilizes its structure after 150-200 ns in all the simulations as shown from the RMSD plots in figures S1, S10, and S19. Only in a few cases does the protein undergo significant structural rearrangements. Most of these rearrangements are related to opening and closing of loop 1 of Smlt1473 (Figure 1B) as highlighted again in figures S1, S10, and S19. In our previous studies, we have discovered that strategies to engineer the flexible loop region could be developed for a wide variety of applications as it is important for the substrate binding^8, 11^. Our current results are complementary to our previous work in highlighting the importance of this loop region.

Next, we studied the importance of fluctuations in the simulation trajectory. We again found that the fluctuations in the flexible loop are the most important in determining the fluctuations in the overall structure (Figures 2, 9, S3, S4, S12, S13, S21, and S22) as well as fluctuations in the distances between the various catalytic residues (Figures S5, S14, and S23).

We found that the overall binding energies of the WT protein for HA_2_, HA_4_, and ManA_6_ sugars are −505±1, −747±2, and −840 ±2 (KJ/mol), respectively. For the H221F mutant, the binding energies are −520±1, −750±2, and −779 ± 2 (KJ/mol), respectively for the sugars. For the R312L mutant the binding energy is −666±1 KJ/mol for ManA. Essentially from these values alone it is hard to draw some correlations about the overall activities. On looking at further per residue interactions, we find that the per residue interactions might be important in determining enzymatic activity. The H221F mutant causes a significant decrease in the interaction of R218 with HA as compared to the WT. On the other hand, H221F causes an increase in the interaction of R218 with the ManA sugar as compared to WT. R218 is an important residue which was previously found to alter the enzyme substrate catalysis for Smlt1473^7, 14^. This might be the reason why H221F causes a decrease in activity for HA as compared to an increase in activity for ManA ^11^ (see Tables 1,2, and 3, and Figures 6, 9, and 12). This effect is caused due to R218 interacting less strongly with the HA sugar (this in turn causes several other residues to interact more strongly with the sugar thereby decreasing the binding energy) as opposed to R218 interacting more strongly with the ManA sugar (this in turn causes several other residues to interact less weakly thereby decreasing the binding energy). For the R312L mutant, there is a decrease in the overall binding energy due to disruption of the R312 salt bridge interacting with the sugar.

To classify the bound states of the three sugars, we performed clustering on these structures to identify stable states in our simulation trajectory. Henceforth, we show the applicability of KMeans and deep learning-based GMVAE in clustering protein-ligand complexes. In Figure 4, S4, S11, S12, S18, and S19, we have shown the various clustering results and have developed a qualitative comparison between K-Means and GMVAE clustering algorithms (see text within those figures). We see that the GMVAE clusters are more physically meaningful in general when compared to the protein-ligand distance plots. This is expected as K-Means depends on a simple Euclidean distance metric as compared to GMVAE which does not rely on an such metric and tries to reconstruct meaningful mixture of Gaussians in the latent space to cluster the data^33^.

The abovementioned clustering was used to provide occupation estimates for the bound states in our MD simulations (Tables S1, S6, and S10). From these individual clusters, we have determined the average binding enthalpies of these clusters (Tables S1, S6, and S9). We note some of the most important values here (from our GMVAE clustering): For HA_2_, cluster 7 is most stable (25.0%) with a binding energy of −483± 1 KJ/mol(WT), cluster 2 was most stable (26.7%) with a binding energy of −562 KJ/mol (H221F). For HA_4_, cluster 1 was most stable (41.1%) with a binding energy of −865± 2 KJ/mol (WT), cluster 3 was most stable (46.1%) with a binding energy of −885± 2 KJ/mol (H221F). For ManA_6_, cluster 2 is most stable (58.9%) with a binding energy of −927 ± 2 KJ/mol (WT), cluster 4 is most stable (42.7%) with a binding energy of −970 ±3 KJ/mol (H221F), cluster 3 is most stable (44.7%) with a binding energy of −703 ± 2 KJ/mol (R312L). The major interactions that stabilize these clusters are mainly salt bridges and hydrogen bonds between the protein and the sugar (Figures 4, 5, 7, 8, S8, S9, S18, and S27, Tables S2-S5, S7, S8, S11-S13). Specifically, R164 and R218 dominate these interactions in HA_4_ for both WT and H221F mutant. In HA_2_, R218 and R312 show important interactions for the WT. For H221F mutant, Y222 and R312 show major interactions with HA_2_. For ManA_6_, R163, R164, and R218 are the residues that show major interactions.

To highlight the ability of our simulations to predict residues that are important for sugar binding and thus the design of better enzymes, this study also included experiments to explore how mutations to newly predicted residues affected Smlt1473 activity. Figures 4, 7, and 10 show the residues that were found to be most important for sugar binding, with K56, R107, and R164 adding to what has already been published^7, 11, 14, 38^. It was found that mutating arginine to glycine at position 107 altered activity against ManA and HA, albeit to different extents. Additionally, mutating lysine to alanine at position 56 altered activity against only HA. Mutating arginine to glycine at position 164 had no effect on activity against ManA or HA. These data show that the simulations were able to predict residues (R107 and K56) that did in fact play a role in sugar binding, as mutations at these positions led to altered activity against ManA and HA. While mutating arginine to glycine at position 164 (R164G) did not result in altered activity for either ManA or HA, this does not necessarily negate its importance in sugar binding. It is perfectly possible that mutating arginine to an amino acid other than glycine at this position could affect binding to these two sugars. Additionally, it is possible that other effects within the protein could be in place that nullify the R164G mutation. To further improve and expand the model, simulating how specific amino acid changes at these already predicted positions affect enzyme activity could further inform better enzyme design with decreased experimental burden. Overall, this study is a start to our future attempts to build a machine-learning model to predict the activity of the Smlt1473 and thus design better enzymes for sustainable products and a better future.

## Acknowledgments

In part, funding for this work supported BWR and SF from an USDA grant #2022-70006-37991. The authors acknowledge the University of Maryland supercomputing resources (http://hpcc.umd.edu) Zaratan made available for conducting the research reported in this paper. This work used Expanse at San Diego Supercomputing Centre through allocation MCB100139 from the <u>Advanced Cyberinfrastructure Coordination Ecosystem: Services &</u> <u>Support</u>(ACCESS) program^39^, which is supported by National Science Foundation grants #2138259, #2138286, #2138307, #2137603, and #2138296.

## Notes

### Competing Interest Statement

The authors have declared no competing interest.

