## Supplementary data for "Effect of Mutations on Smlt1473 Binding to Various Substrates Using Molecular Dynamics Simulations"

**Table S1**: Binding energies for the clusters for sugar $\mathrm{HA}_{2}$ predicted by our KMeans and GMVAE algorithm for both the wild-type as well as the H221F mutation. Also noted are the total fraction present for each of the clusters.

| **Cluster No** | **WT KMeans Fraction** | **WT**  **KMeans**  **(in**  **KJ/mol)** | **H221F**  **KMeans**  **Fraction** | **H221F**  **KMeans**  **(in**  **KJ/mol)** | **GMVAE**  **WT**  **Fraction** | **WT**  **GMVAE**  **(in**  **KJ/mol)** | **GMVAEH221F**  **fraction** | **H221F**  **GMVAE**  **(in**  **KJ/mol)** |
| --- | --- | --- | --- | --- | --- | --- | --- | --- |
| 0 | 0.01% | -311 | 6.51% | -448±2 | 6.71% | -409±3 | 7.57% | -46±3 |
| 1 | 0.00% | NAN | 2.26% | -200$\pm$3 | 0.65% | -483$\pm$13 | 16.79% | -490$\pm$2 |
| 2 | 8.05% | -501$\pm$ 3 | 56.53% | -525$\pm$1 | 0.90% | -506$\pm$9 |  | -562$\pm$1.50 |
| 3 | 16.36% | -433$\pm$2 | 0.61% | -460$\pm$8 | 15.80% | -470$\pm$3 | 1.53% | -504$\pm$6 |
| 4 | 0.00% |  | 2.49% | -297$\pm$3 | 2.61% | -488$\pm$6 | 4.53% | -551$\pm$5 |
| 5 | 1.97% | -431$\pm$6 | 20.76% | -632$\pm$2 | 0.45% | -524$\pm$12 | 19.87% | -636$\pm$2 |
| 6 | 19.25% | -684$\pm$2 |  |  | 15.42% | -722$\pm$2 |  |  |
| 7 | 23.84% | -475$\pm$1 |  |  | 25.01% | -483$\pm$1 |  |  |
| 8 | 4.20% | -423$\pm$4 | 0.06% | -435$\pm$33 | 7.86% | -477$\pm$3 | 4.62% | -480$\pm$2 |
| 9 | 11.11% | -547$\pm$ 2 | 4.61% | -473$\pm$3 | 10.11% | -529±3 | 0.96% | -504$\pm$12 |
| 10 | 2.12% | -499$\pm$ 5 | 6.18% | -441$\pm$ 2 | 2.10% | -470$\pm$6 | 6.24% | -440$\pm$2 |
| 11 | 13.09% | -406 $\pm$ 2 | 0.01% | -299 | 12.38% | -400$\pm$2 | 11.21% | -365$\pm$3 |

**Table S2**: A table showing the interaction energies of the 10 most highly interacting residues from the WT simulation data from our HA_2_ simulation for the maximal populate clusters.

| Residue | Cluster 3  (in KJ/mol) | Cluster 6  (in KJ/mol) | Cluster 7  (in KJ/mol) | Cluster 8  (in KJ/mol) | Cluster 9  (in KJ/mol) | Cluster 11  (in KJ/mol) |
| --- | --- | --- | --- | --- | --- | --- |
| Y38 | -37.5 | -57.3 | -28.8 | -19.6 | -11.9 | -29.4 |
| Y39 | -40.2 | -67.9 | -74.6 | -10.1 | -18.3 | -7.05 |
| K56 | -31.6 | -1.04 | -22.5 | -0.00497 | -0.319 | -3.50 |
| Q112 | -18.8 | -38.7 | -42.2 | -2.60 | -4.44 | -0.304 |
| H168 | -13.7 | -30.8 | -36.6 | -18.5 | -19.3 | -1.51 |
| R215 | -19.2 | -14.4 | -3.39 | -132 | -74.9 | -3.75 |
| R218 | -71.9 | -89.7 | -161 | -92.5 | -82.1 | -107 |
| Y222 | -38.8 | -39.1 | -26.1 | -39.2 | -50.2 | -8.37 |
| Y225 | -9.33 | -41.2 | -11.3 | -19.3 | -20.4 | -10.1 |
| R312 | -89.0 | -163 | -2.76 | -2.35 | -150 | -147 |

**Table S3**: A table showing the interaction energies of the 10 most highly interacting residues from the H221F simulation data from our HA_2_ simulation for the maximal populated clusters.

| Residue | Cluster 0  (in KJ/mol) | Cluster 1  (in KJ/mol) | Cluster 2  (in KJ/mol) | Cluster 5  (in KJ/mol) | Cluster 10  (in KJ/mol) | Cluster 11  (in KJ/mol) |
| --- | --- | --- | --- | --- | --- | --- |
| G37 | -1.37 | -21.9 | -24.8 | -1.61 | -0.142 | -0.0253 |
| Y38 | -28.8 | -45.6 | -48.4 | -31.2 | -10.8 | -18.6 |
| Y39 | -21.4 | -41.8 | -55.3 | -36.6 | -23.8 | -24.5 |
| K42 | -7.30 | -25.2 | -31.9 | -0.115 | 0.466 | -0.104 |
| Q112 | -29.0 | -27.8 | -32.8 | -22.1 | -0.674 | -0.100 |
| R215 | -21.4 | -23.4 | -26.7 | -168 | -112 | -5.98 |
| R218 | -9.20 | -10.6 | -10.4 | -33.7 | -165 | -165 |
| Y222 | -59.3 | -63.5 | -58.4 | -8.29 | -12.9 | -9.00 |
| Y225 | -46.7 | -44.6 | -44.3 | -62.5 | -0.171 | -1.32 |
| R312 | -62.2 | -92.0 | -114 | -162 | -0.0107 | -0.746 |

**Table S4**: A table showing the interaction energies of the 10 most highly interacting residues in the WT simulation data from the less populated clusters for sugar $\mathrm{HA}_{2}$.

| Residue | Cluster 0  (KJ/mol) | Cluster 1  (KJ/mol) | Cluster 2  (KJ/mol) | Cluster 4  (KJ/mol) | Cluster 5  (KJ/mol) | Cluster 10  (KJ/mol) |
| --- | --- | --- | --- | --- | --- | --- |
| Y38 | -34.2 | -41.2 | -47.4 | -45.0 | -23.5 | -21.5 |
| Y39 | -28.6 | -56.8 | -59.7 | -62.1 | -36.0 | -40.7 |
| K56 | -1.15 | -1.00 | -14.2 | -11.4 | -2.85 | -8.69 |
| Q112 | -2.08 | -21.7 | -27.7 | -21.0 | -23.4 | -6.88 |
| H168 | -25.7 | -2.43 | 2.20 | 1.28 | -11.0 | -4.47 |
| R215 | -18.2 | -13.5 | -10.7 | -24.5 | -148 | -101 |
| R218 | -71.5 | -92.0 | -78.4 | -96.8 | -0.900 | -159 |
| Y222 | -39.1 | -55.9 | -49.6 | -36.4 | -8.59 | -21.2 |
| Y225 | -35.4 | -23.9 | -18.3 | -12.8 | -1.46 | -2.78 |
| R312 | -69.3 | -60.6 | -63.4 | -55.6 | -168 | -0.0344 |

**Table S5**: A table showing the interaction energies of the 10 most highly interacting residues in the H221F simulation data from the less populated clusters for sugar $\mathrm{HA}_{2}$.

| Residue | Cluster 3  (in KJ/mol) | Cluster 4  (in KJ/mol) | Cluster 8  (in KJ/mol) | Cluster 9  (in KJ/mol) |
| --- | --- | --- | --- | --- |
| G37 | -23.9 | -25.0 | -1.16 | -0.758 |
| Y38 | -52.6 | -52.2 | -20.9 | -27.7 |
| Y39 | -56.3 | -59.3 | -25.1 | -22.9 |
| K42 | -30.4 | -30.5 | -0.174 | -0.491 |
| Q112 | -33.9 | -32.7 | -17.0 | -15.2 |
| R215 | -19.7 | -32.5 | -150 | -167 |
| R218 | -6.58 | -9.84 | -45.6 | -27.2 |
| Y222 | -31.8 | -54.4 | -72.8 | -14.2 |
| Y225 | -5.56 | -11.8 | -11.3 | -44.0 |
| R312 | -54.5 | -80.0 | -4.27 | -94.8 |

**Table S6**: A table showing the various binding energies in the various clusters predicted by our KMeans and GMVAE algorithm for both the wild-type as well as the H221F mutation for sugar $\mathrm{HA}_{4}$. We also note the total fraction present for each of the clusters.

| Cluster No | WT  KMeans Fraction | WT  KMeans  (in KJ/mol) | H221F  KMeans  Fraction | H221F  (KMeans) | GMVAE  WT  Fraction | GMVAE  WT  (in KJ/mol) | H221F  GMVAE  fraction | H221F (GMVAE)  (in  KJ/mol) |
| --- | --- | --- | --- | --- | --- | --- | --- | --- |
| 0 | 46.85% | -736$\pm$2 | 6.41% | -662$\pm$4 | 31.94% | -687$\pm$2 | 17.35% | -602$\pm$3 |
| 1 | 25.60% | -877$\pm$3 | 63.92% | -856$\pm$1 | 41.13% | -865$\pm$2 | 15.19% | -790$\pm$3 |
| 2 | 25.72% | -657$\pm$3 | 14.75% | -541$\pm$3 | 26.93% | -647$\pm$3 | 21.40% | -564$\pm$3 |
| 3 | 1.84% | -595$\pm11$ | 14.92% | -557$\pm$3 |  |  | 46.06% | -885$\pm2$ |

**Table S7**: A table showing the interactions energies of the 10 most highly interacting residues with the sugar $\mathrm{HA}_{4}$ for the wild type simulation

| Residue | Cluster 0 (in KJ/mol) | Cluster 1  (in KJ/mol) | Cluster 2  (in KJ/mol) |
| --- | --- | --- | --- |
| Y38 | -28.6 | -48.5 | -28.9 |
| Y39 | -29.8 | -47.6 | -24.5 |
| R107 | -19.8 | -47.8 | 0.192 |
| Y115 | -32.5 | -66.3 | -0.826 |
| R163 | -96.1 | -136 | -3.153 |
| R164 | -73.3 | -25.8 | -90.3 |
| R215 | -52.8 | -24.3 | -98.6 |
| R218 | -168 | -91.3 | -201 |
| Y222 | -25.2 | -54.8 | -12.5 |
| Y225 | -7.55 | -38.0 | -8.74 |

**Table S8**: A table showing the interactions energies of the 10 most highly interacting residues with the sugar $\mathrm{HA}_{4}$ for the H221F-mutant simulation.

| Residue | Cluster 0  (in KJ/mol) | Cluster 1  (in KJ/mol) | Cluster 2  (in KJ/mol) | Cluster 3  (in KJ/mol) |
| --- | --- | --- | --- | --- |
| Y38 | -10.3 | -54.2 | -22.5 | -77.5 |
| Y39 | -28.6 | -52.0 | -24.6 | -22.4 |
| K42 | -80.6 | -19.8 | -31.5 | -41.6 |
| R107 | -0.900 | -36.0 | -7.98 | -142 |
| R163 | -9.62 | -125 | -1.88 | -12.2 |
| R164 | -87.1 | -45.5 | -82.4 | -71.1 |
| R215 | -79.4 | -25.8 | -92.7 | -124 |
| R218 | -178 | -68.8 | -95.9 | -28.1 |
| Y222 | -19.1 | -60.6 | -27.4 | -40.8 |
| R312 | -0.295 | -23.6 | -22.0 | -54.4 |

**Table S9**: A table showing the various binding energies in the various clusters predicted by our KMeans and GMVAE algorithm for both the wild-type, H221F, and R312L mutation for the sugar $\mathrm{MANA}_{6}$. We also note the total fraction present for each of the clusters.

| **Cluster No** | **WT**  **KMeans**  **(in**  **KJ/mol)** | **H221F**  **KMeans**  **(in**  **KJ/mol)** | **R312L**  **KMeans**  **(in**  **KJ/mol)** | **WT**  **GMVAE**  **(in**  **KJ/mol)** | **H221F**  **GMVAE**  **(in**  **KJ/mol)** | **R312L**  **GMVAE**  **(in**  **KJ/mol)** |
| --- | --- | --- | --- | --- | --- | --- |
| 0 | -500 $\pm$ 3 | -533 $\pm$ 2 |  | -710 $\pm$ 2 | -496 $\pm$ 3 |  |
| 1 |  | -272 $\pm$ 12 | -586 $\pm$ 2 | -371 $\pm$ 33 | -572 $\pm$ 2 | -585 $\pm$ 2 |
| 2 | -915 $\pm$ 2 | -919 $\pm$ 5 | -702 $\pm$ 3 | -927 $\pm$ 2 | -774 $\pm$ 5 | -604 $\pm$ 4 |
| 3 |  | -626 $\pm$ 4 | -700 $\pm$ 2 |  | -631$\pm$ 4 | -703 $\pm$ 2 |
| 4 | -789 $\pm$ 2 | -428 $\pm$ 3 |  | -761 $\pm$ 7 | -970 $\pm$3 | -765 $\pm$ 3 |

**Table S10**: A table showing the various fractions in the various clusters predicted by our KMeans and GMVAE algorithm for both the wild-type, H221F, and R312L mutation for the sugar $\mathrm{MANA}_{6}$.

| **Cluster**  **No** | **Fraction**  **KMeans**  **WT**  **(in**  **KJ/mol)** | **Fraction**  **KMeans**  **H221F**  **(in**  **KJ/mol)** | **Fraction**  **KMeans**  **R312L**  **(in**  **KJ/mol)** | **Fraction**  **GMVAE**  **WT**  **(in**  **KJ/mol)** | **Fraction**  **GMVAE**  **H221F**  **(in**  **KJ/mol)** | **Fraction**  **GMVAE**  **R312L**  **(in**  **KJ/mol)** |
| --- | --- | --- | --- | --- | --- | --- |
| 0 | 10.08% | 19.80% | 0.00% | 36.80% | 12.17% | 0.00% |
| 1 | 0.00% | 0.43% | 30.20% | 0.11% | 13.47% | 30.54% |
| 2 | 63.18% | 59.38% | 20.38% | 58.87% | 19.56% | 10.01% |
| 3 | 0.00% | 20.04% | 49.42% | 0.00% | 12.16% | 44.67% |
| 4 | 26.74% | 0.35% | 0.00% | 4.23% | 42.65% | 14.78% |

**Table S11**: A table showing the interaction energies of the most important residues to the sugar in each of the individual clusters for the Wild-Type simulation for the sugar $\mathrm{MANA}_{6}$.

| Residue | Cluster 0  (in KJ/mol) | Cluster 1  (in KJ/mol) | Cluster 2  (in KJ/mol) | Cluster 4  (in KJ/mol) |
| --- | --- | --- | --- | --- |
| K42 | -99.4 | -39.0 | -27.8 | -61.1 |
| K56 | 0 | 0 | -113 | -22.5 |
| R107 | -137 | -156 | -4.86 | -76.7 |
| K162 | -56.1 | -10.8 | 0 | -2.16 |
| R163 | -162 | -102 | -78.9 | -49.0 |
| R164 | -126 | -8.64 | -122 | -55.1 |
| R215 | -101 | -12.3 | -148 | -108 |
| R218 | 0 | -33.8 | -137 | -112 |
| Y222 | 0 | 0 | -60.4 | -25.4 |
| R312 | 0 | 0 | -124 | -73.0 |

**Table S12**: A table showing the interaction energies of the most important residues to the sugar in each of the individual clusters for the H221F mutant simulation for the sugar $\mathrm{MANA}_{6}$.

| Residue | Cluster 0  (in KJ/mol) | Cluster 1  (in KJ/mol) | Cluster 2  (in KJ/mol) | Cluster 3  (in KJ/mol) | Cluster 4  (in KJ/mol) |
| --- | --- | --- | --- | --- | --- |
| K42 | -4.23 | -0.831 | -2.03 | -42.8 | -68.5 |
| K56 | 0 | 0 | -118 | -23.0 | -33.4 |
| R107 | -172 | -157 | 0 | -57.2 | -12.1 |
| K162 | -123 | -89.7 | -0.483 | -43.4 | -16.1 |
| R163 | -159 | -156 | -3.27 | -7.81 | -4.06 |
| R164 | -5.94 | -139 | -87.5 | -133 | -119 |
| R215 | 0 | -5.85 | -137 | -87.4 | -137 |
| R218 | 0 | -1.40 | -134 | -121 | -168 |
| Y222 | 0 | -0 | -50.3 | -23.4 | -58.9 |
| R312 | 0 | 0 | -62.9 | 0 | -102 |

**Table S13**: A table showing the interaction energies of the most important residues to the sugar for the R312L mutant simulation for the sugar $\mathrm{MANA}_{6}$.

| Residue | Cluster 1  (in KJ/mol) | Cluster 2  (in KJ/mol) | Cluster 3  (in KJ/mol) | Cluster 4  (in KJ/mol) |
| --- | --- | --- | --- | --- |
| K42 | -2.55 | -5.68 | -98.0 | -8.71 |
| K56 | 0 | -58.8 | -0.0328 | -67.6 |
| K162 | -14.94 | 0.159 | -135 | -9.52 |
| R163 | -172 | -27.4 | -12.3 | -105 |
| R164 | -110 | -77.7 | -158 | -150 |
| R165 | -162 | 0.0172 | -0.974 | 0.041 |
| K198 | -112 | 0.000337 | -0.0665 | -0.00357 |
| R215 | -0.664 | -83.032 | -109 | -92.6 |
| R218 | -0.00675 | -165 | -113 | -138 |
| Y222 | -0.000168 | -43.8 | -23.1 | -22.9 |

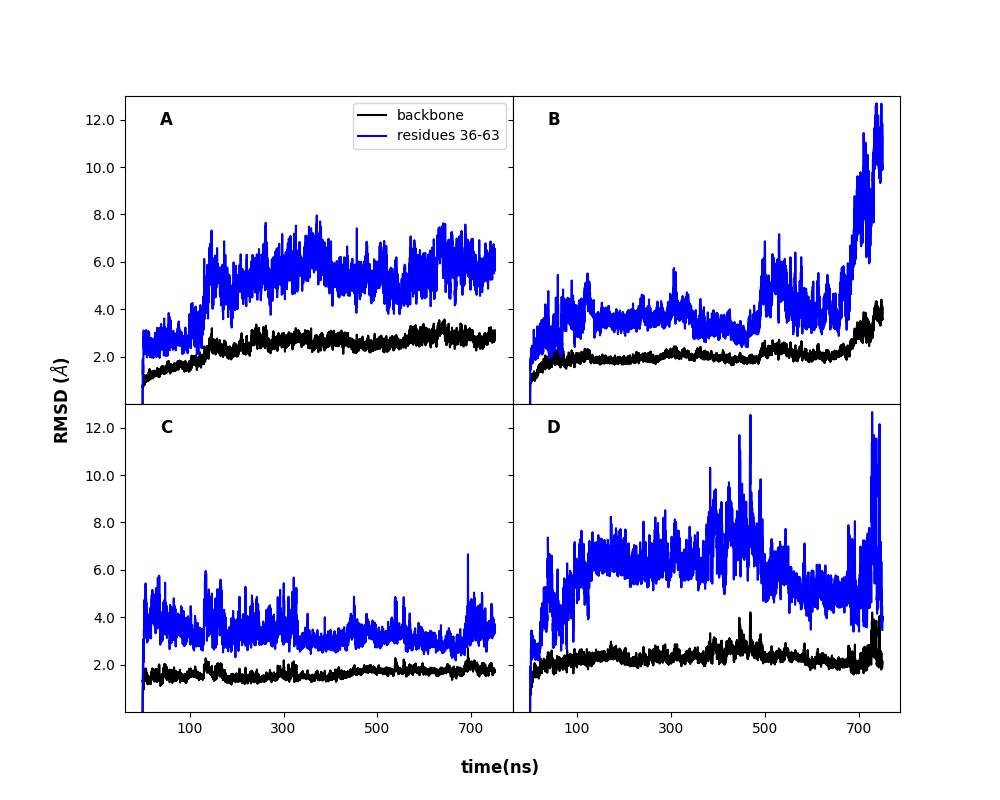

**Figure S1**: RMSD of the protein backbone with HA_2_ as well as the entrance loop with the adjacent helical region (residues 36-63, see Fig. 1B) for WT-set1 (A), WT-set2 (B), H221F-set1 (C), H221F-set2 (D).

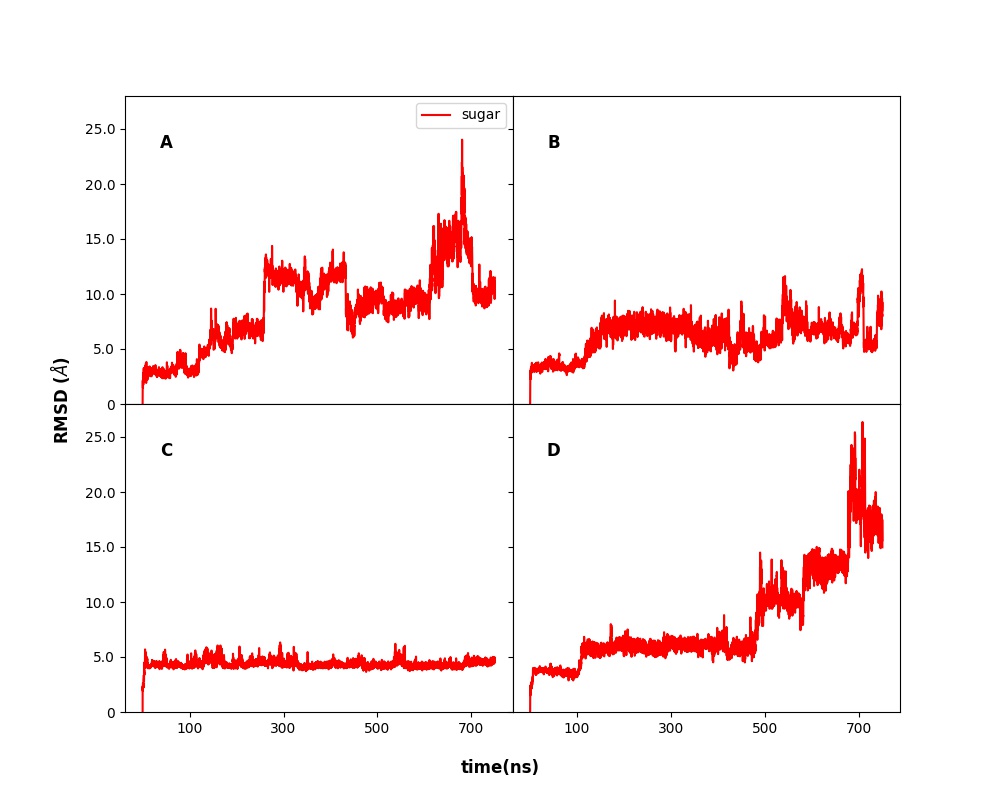

**Figure S2**: RMSD of the sugar $\mathrm{HA}_{2}$ as a function of time

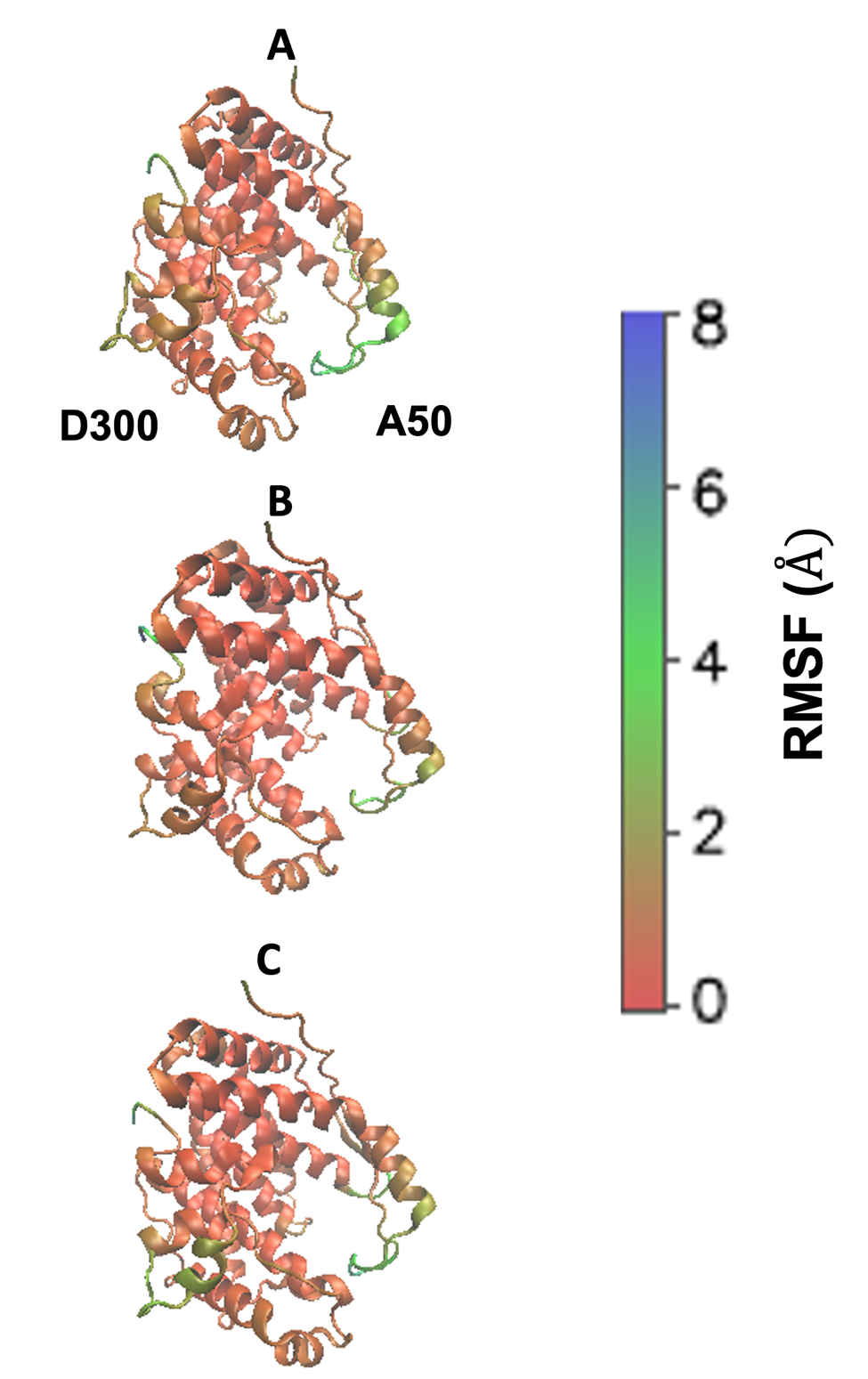

**Figure S3:** RMSF for HA2 replicas WT-set2 (A), H221F-set1 (C), and H221F-set2 (D).

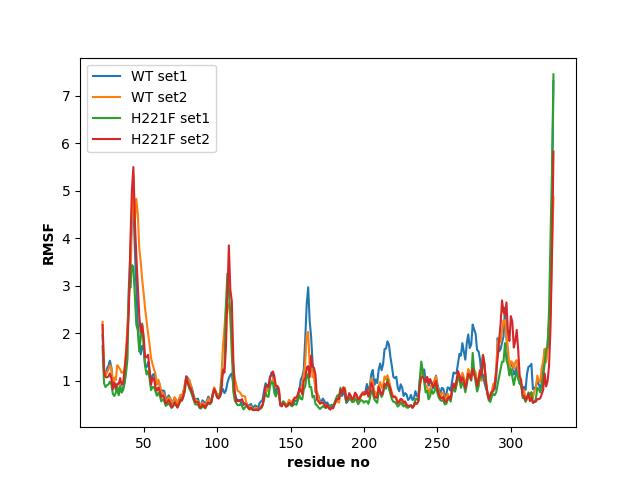

**Figure S4**: The RMSF of the all protein residues from the entire four sets of simulations for sugar $\mathrm{HA}_{2}$ .

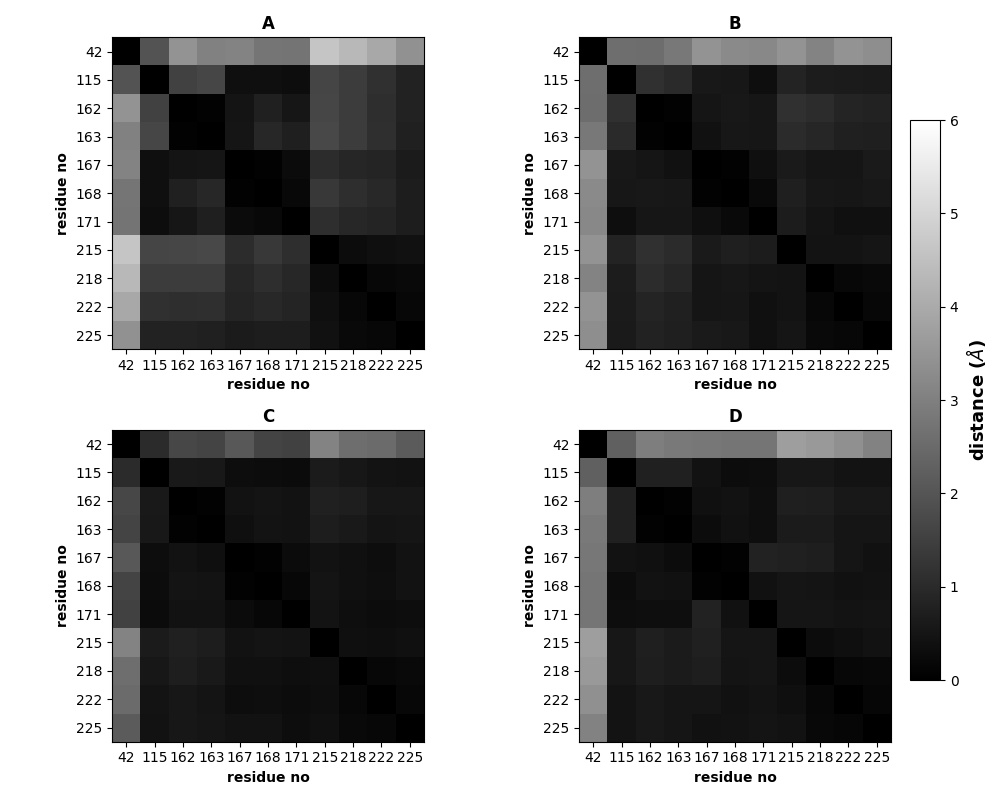

**Figure S5:** A(for WT-set1), B(for WT-set2), C(for H221F-set1), D(for H221F-set2) respectively denotes the various regions on the protein for the sugar $\mathrm{HA}_{2}$ with varying fluctuations.

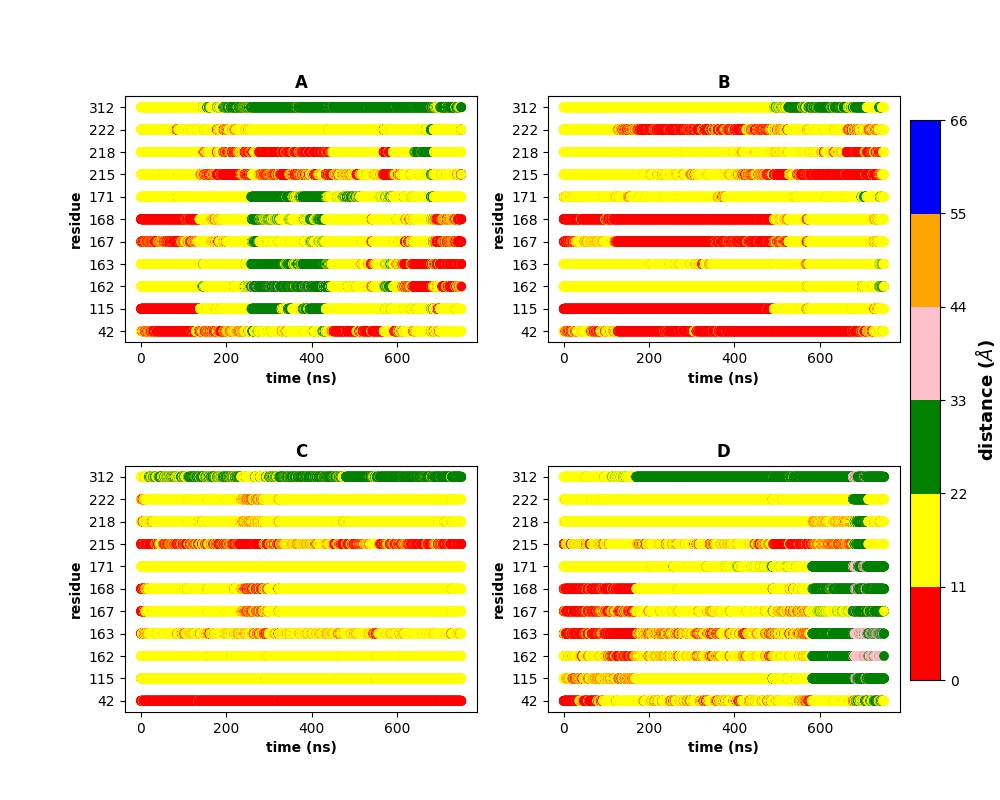

**Figure S6:** (A) and (B) denotes the distances between each of the important residues on the protein to the centre of mass of the sugar for simulations of WT-set1 and set2 respectively. (C) and (D) denotes- the same distances between H221F-mutant and the centre of mass of the sugar respectively for the two sets of HA_2_ simulations, respectively.

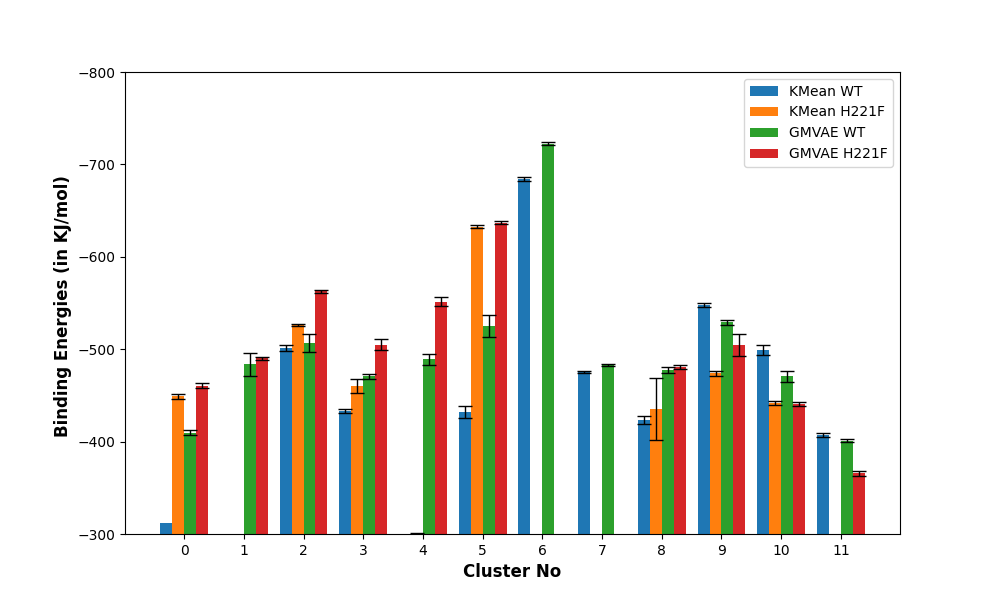

**Figure S7:** The binding energies (±standard error) of the various stable clusters from the various simulation sets with HA_2_. The energy values are not present in some x-axis ranges which means that that particular cluster was absent from the simulations. The detailed data about the various clusters are shown in Table S1.

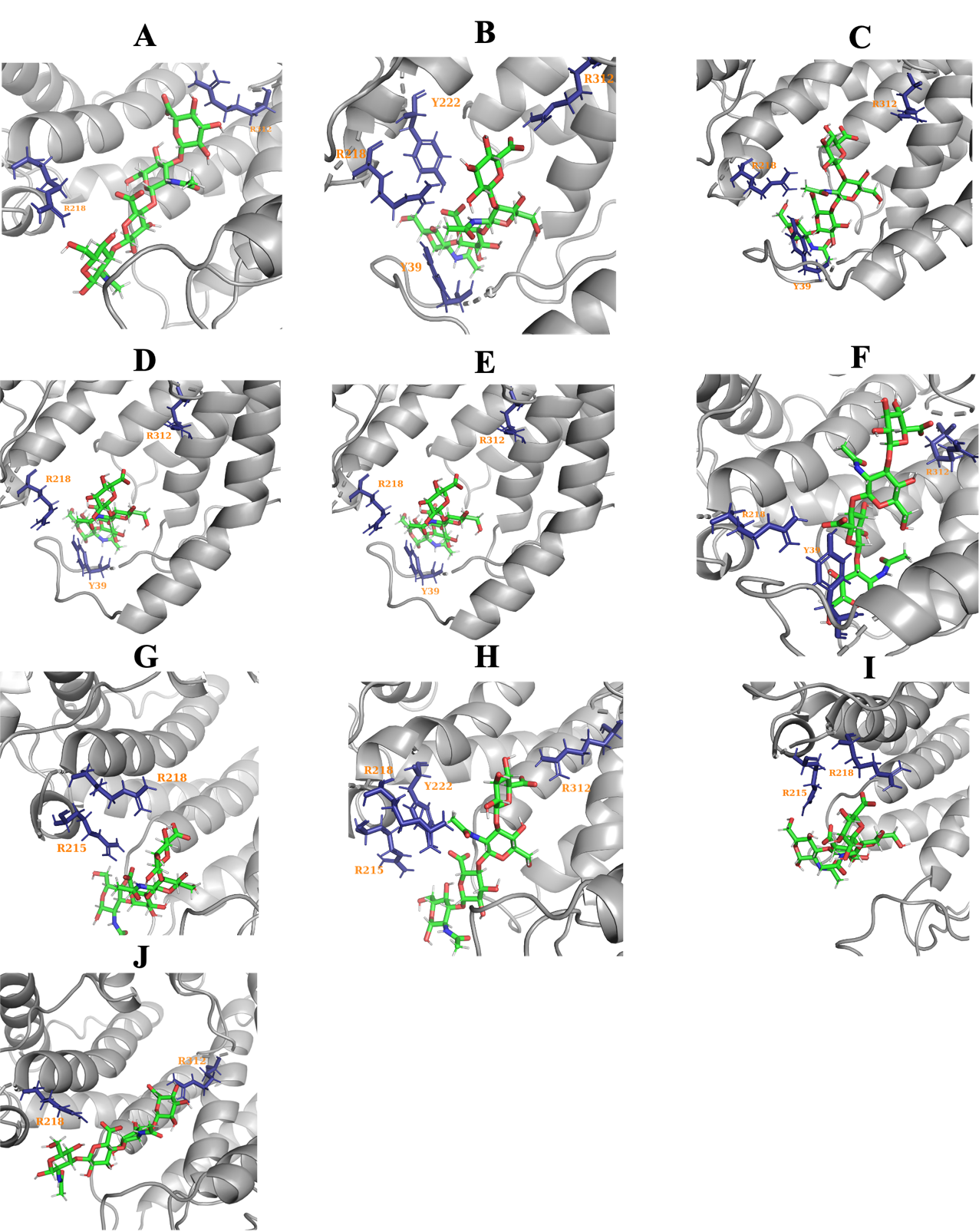
 **Figure S8**: The less populated clusters and their major interactions (see text) predicted by our GMVAE algorithm for the WT simulation with HA_2_.Here are the cluster codes: A- 0, B-1, C-2, D-4, E-5, F-6, G-8, H-9, I-10, J-11.

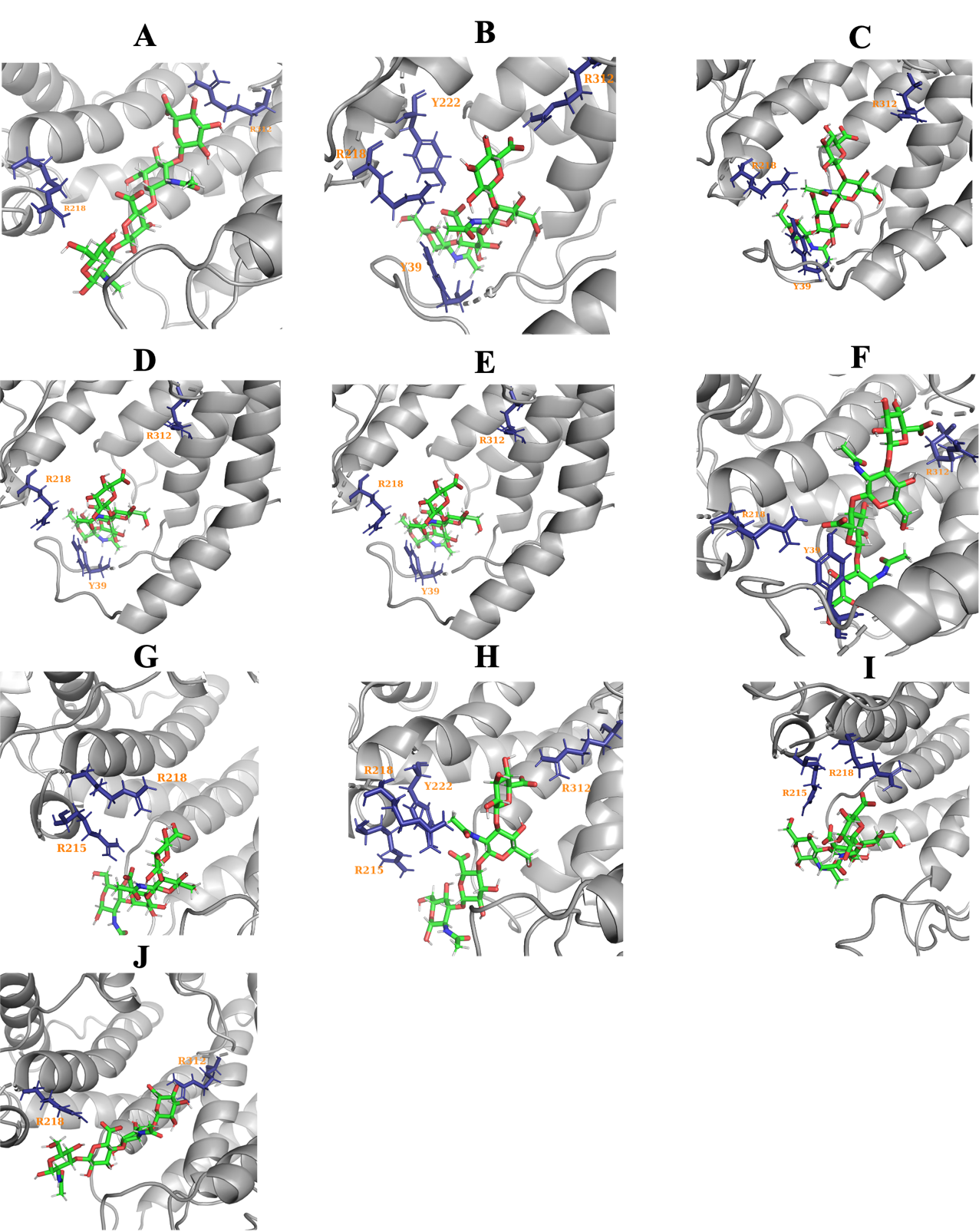

**Figure S9**: The less populated clusters and their major interactions (see text) predicted by our GMVAE algorithm for the H221F simulation with HA_2_.Here are the cluster codes: A- 0, B-1, C-3, D-4, E-8, F-9, G-10, J-11.

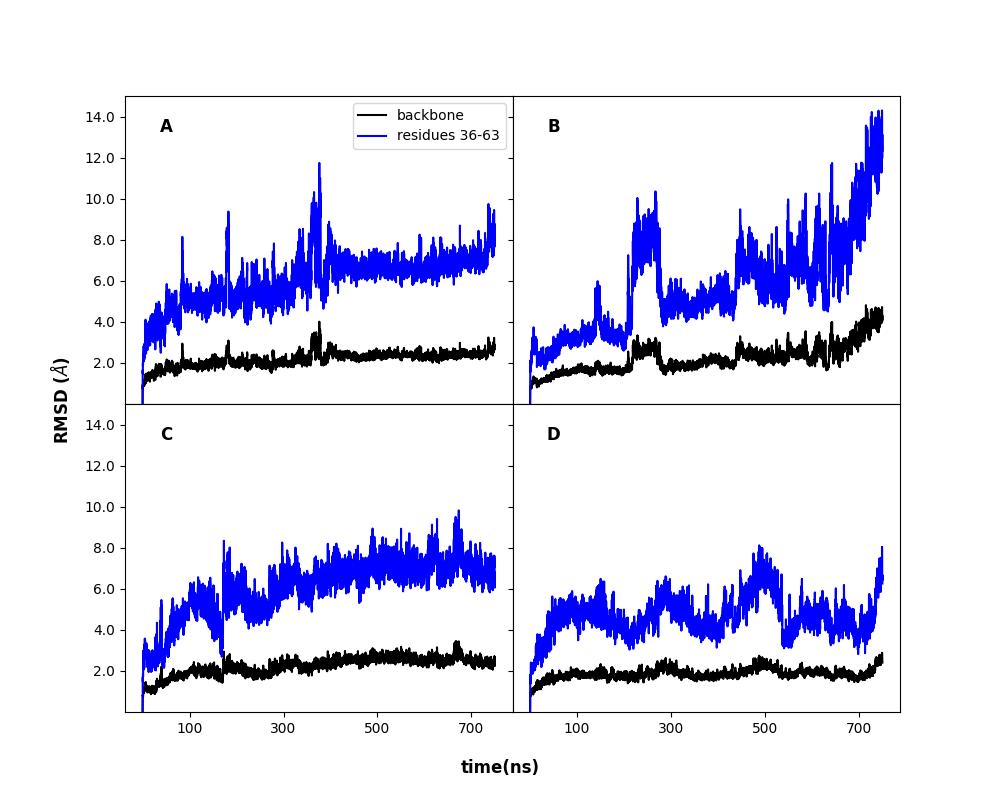

**Figure S10**: RMSD of the protein backbone with HA_4_ as well as the entrance loop with the adjacent helical region (residues 36-63, see Fig. 1B) for A(WT-set1), B(WT-set2), C(H221F-set1), and D(H221F-set2).

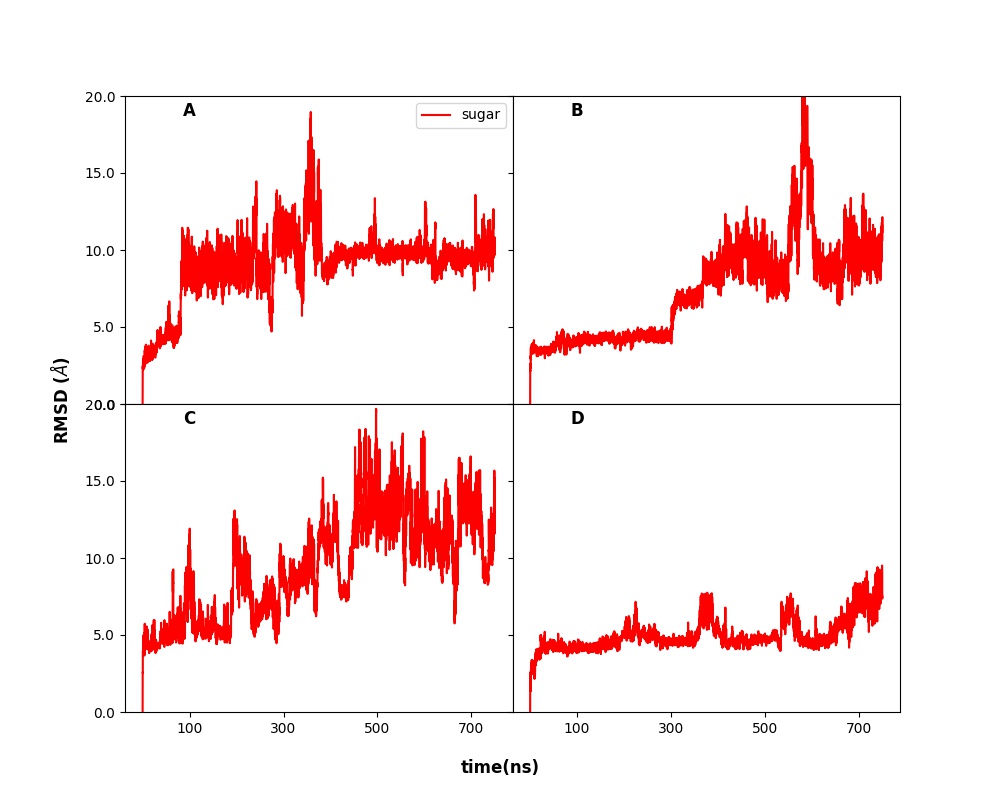

**Figure S11**: RMSD of the sugar $\mathrm{HA}_{4}$ as a function of time

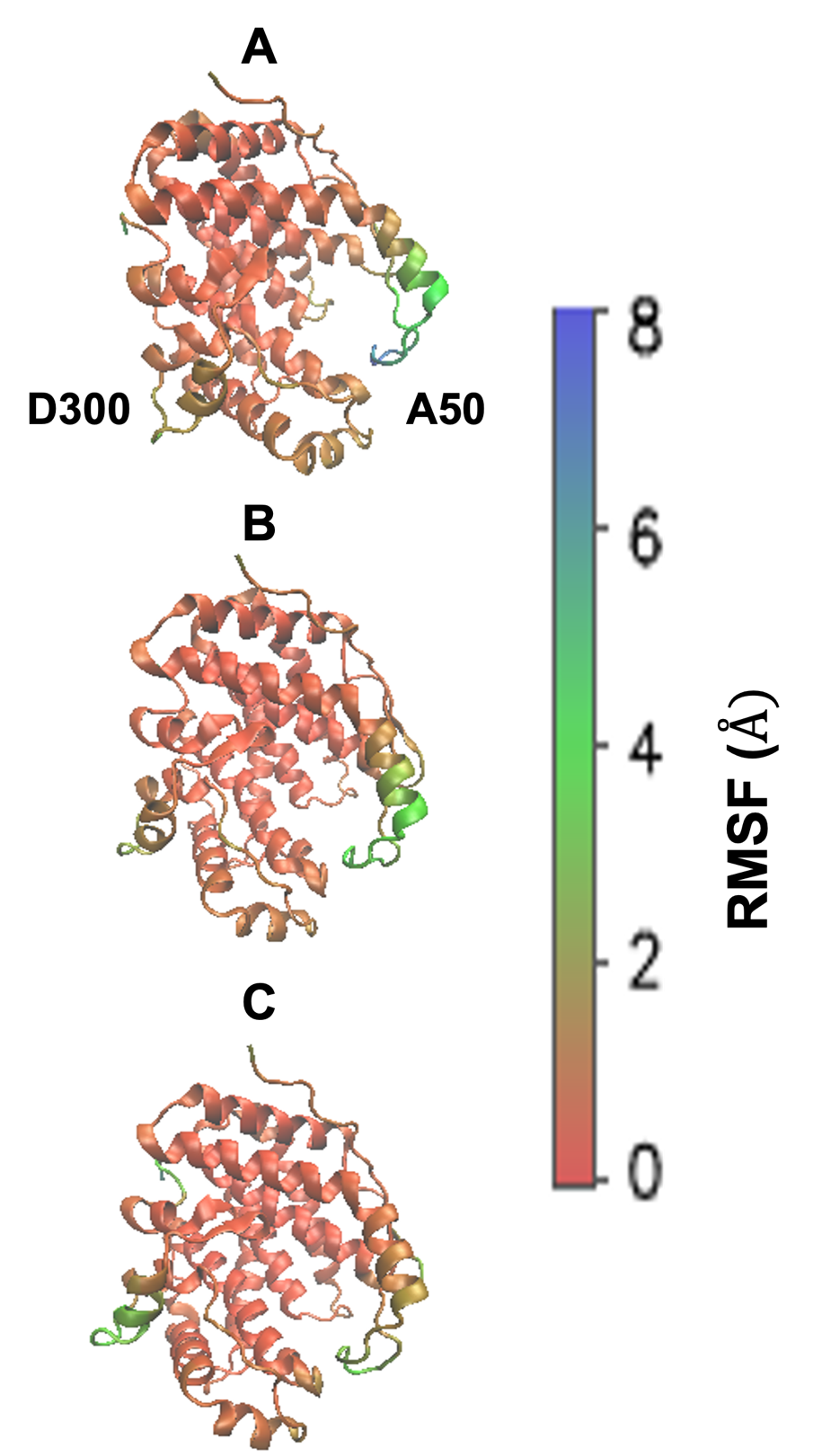

**Figure S12:** RMSF for $\mathrm{HA}_{4}$ replicas for WT-set2 (A), H221F-set1 (C), and H221F-set2 (D).

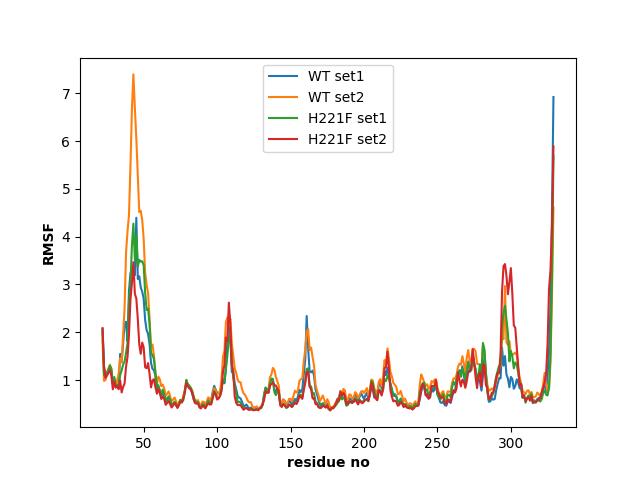

**Figure S13**: A(for WT-set1), B(for WT-set2), C(for H221F-set1), D(for H221F-set2) respectively denote the RMSF of the various residues on the protein for the sugar $\mathrm{HA}_{4}$.

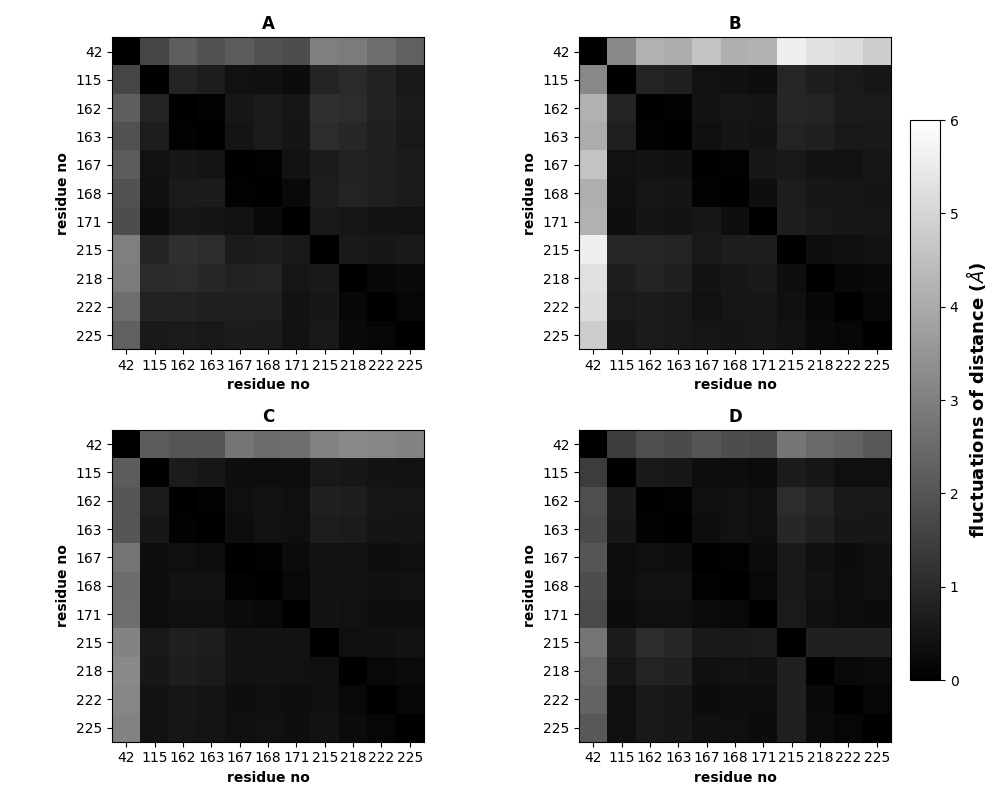

**Figure S14**: (A)-for WT set1, (B) for WT set-2, (C) for H221F- set1, and (D) for H221F- set2 denote the fluctuations of the distances between the various residues of the protein for the sugar $\mathrm{HA}_{4}$.

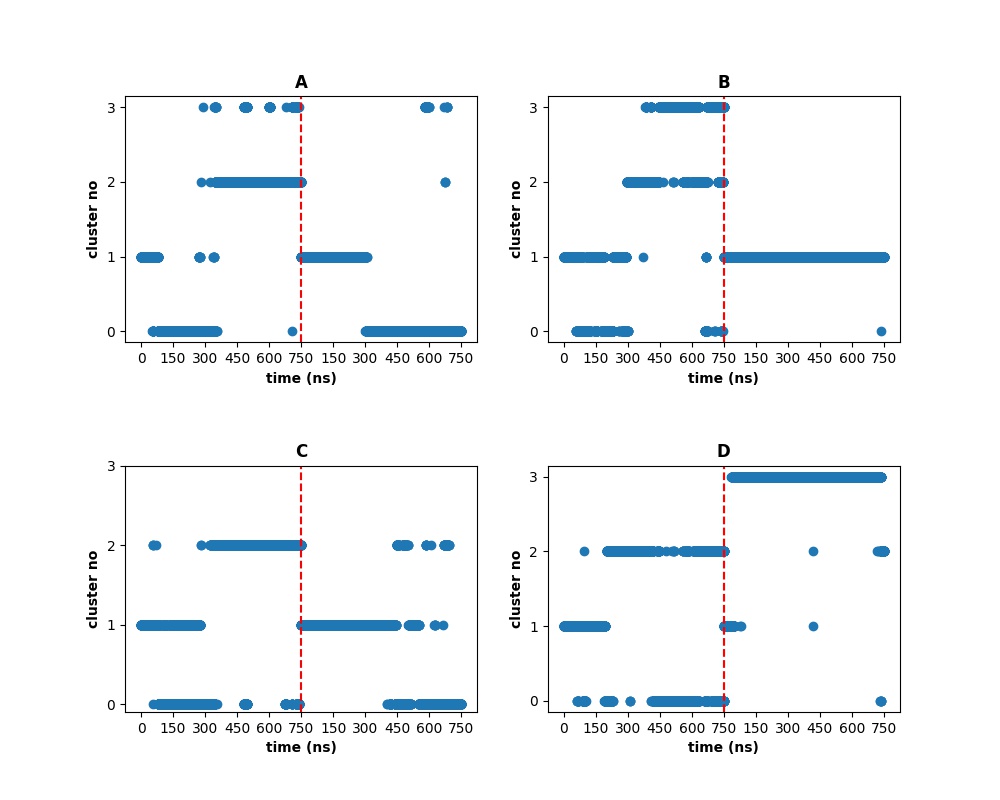

**Figures S15**: (A)-Wild type and (B)-H221F denote the various clusters predicted by the KMeans for the various structures of the protein-sugar ($\mathrm{HA}_{4}$) complex for our sugar $\mathrm{HA}_{4}$. (C)-Wild type and (D)-H221F denotes the various clusters predicted by GMVAE for the various structures for our sugar $\mathrm{HA}_{4}$. The red line in between seperates the two sets for the simulation.

The states have been clustered based on the distances of the centre of masses of the various sugar units to the closest residues on the protein. The GMVAE shows a better clustering of states for HA_4_ as compared to a normal K-Means. For example, the GMVAE shows a much wider variation in energy levels as compared to the K-Means (Table S6 ). Also, the variation in the contact maps (Figure S13) as a function of the simulation time shows more resemblence to the fluctuations in the clusters in the GMVAE case. For example, on observing the $\mathrm{HA}_{4}$ simulation distances in Figure S13B, we see that for 0-400 ns the colors/distances shown in the trajectory do not change and there is a change in distance only after 400 ns. This fact is verified by our GMVAE algorithm right of red line in Figure S12C which shows a change in cluster only after 400 ns but not by the KMeans from Figure S12A. Similarly, for $\mathrm{HA}_{4}$- H221F mutation for set2, we can see that the there is a change in distance as compared to the initial color during the start of the simulation from figure S13D, which is not reflected by K-Means but instead reflected by the GMVAE.

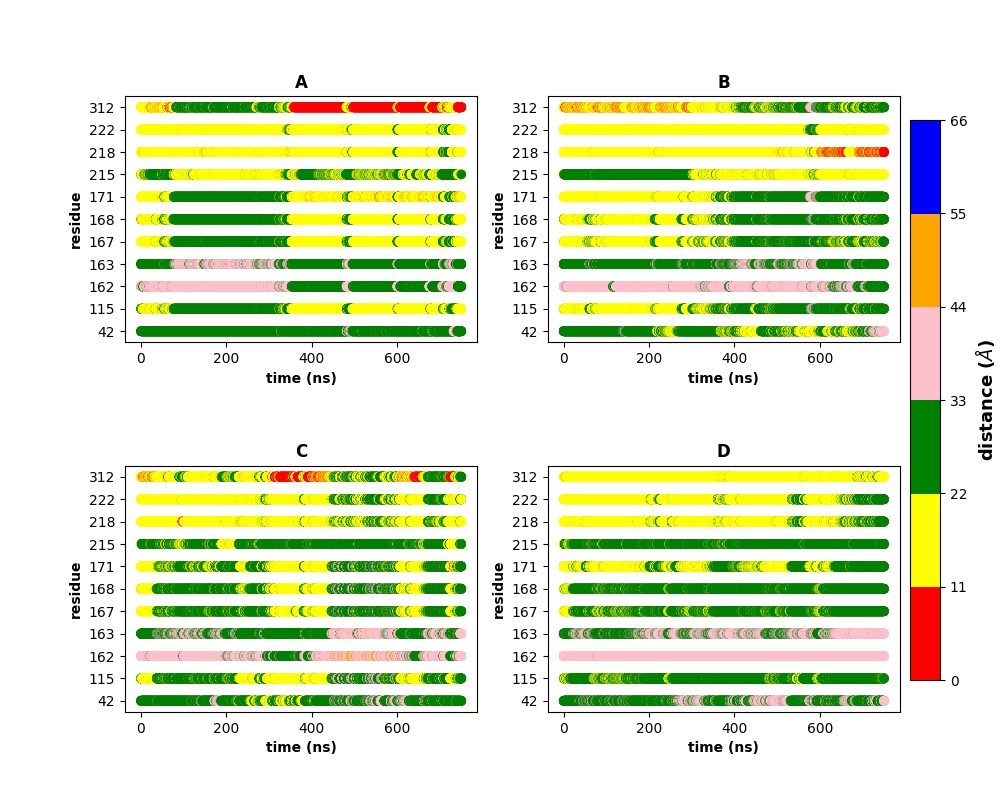
**Figure S16**: (A) and (B) denotes the distances between each of the important residues on the protein to the centre of mass of the sugar $\mathrm{HA}_{4}$ for simulations of wild type-set1 and set2 respectively. (C) and (D) denotes- the same distances between H221F-mutant and the centre of mass of the sugar $\mathrm{HA}_{4}$ respectively for the two sets of simulation, respectively.

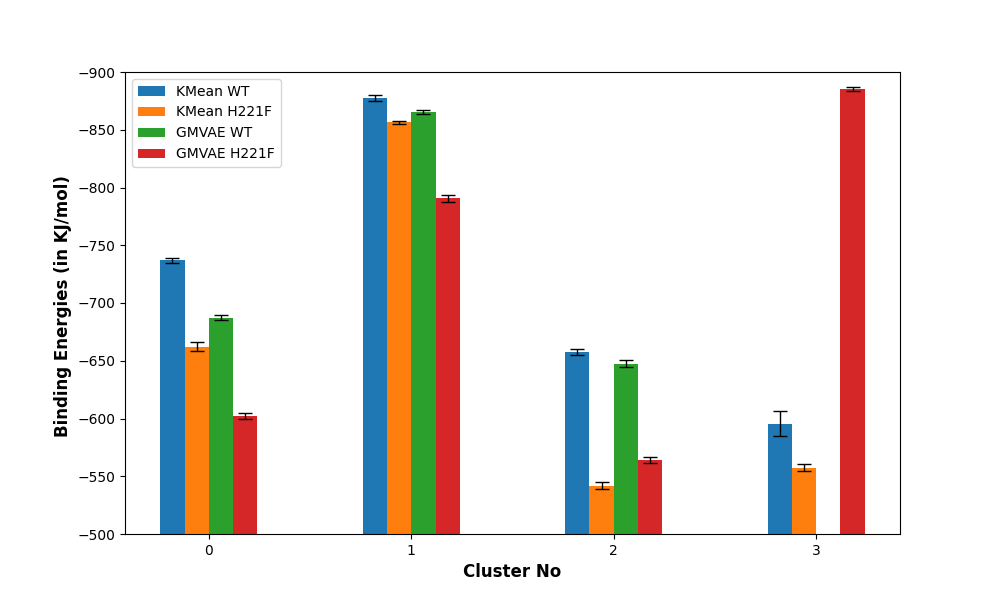

**Figure S17**: The binding energies of the various stable clusters from the various simulation sets with HA_4_. The energy values are not present in some x-axis ranges which means that that particular cluster was absent from the simulations. The detailed data about the various clusters are shown in Table S6.

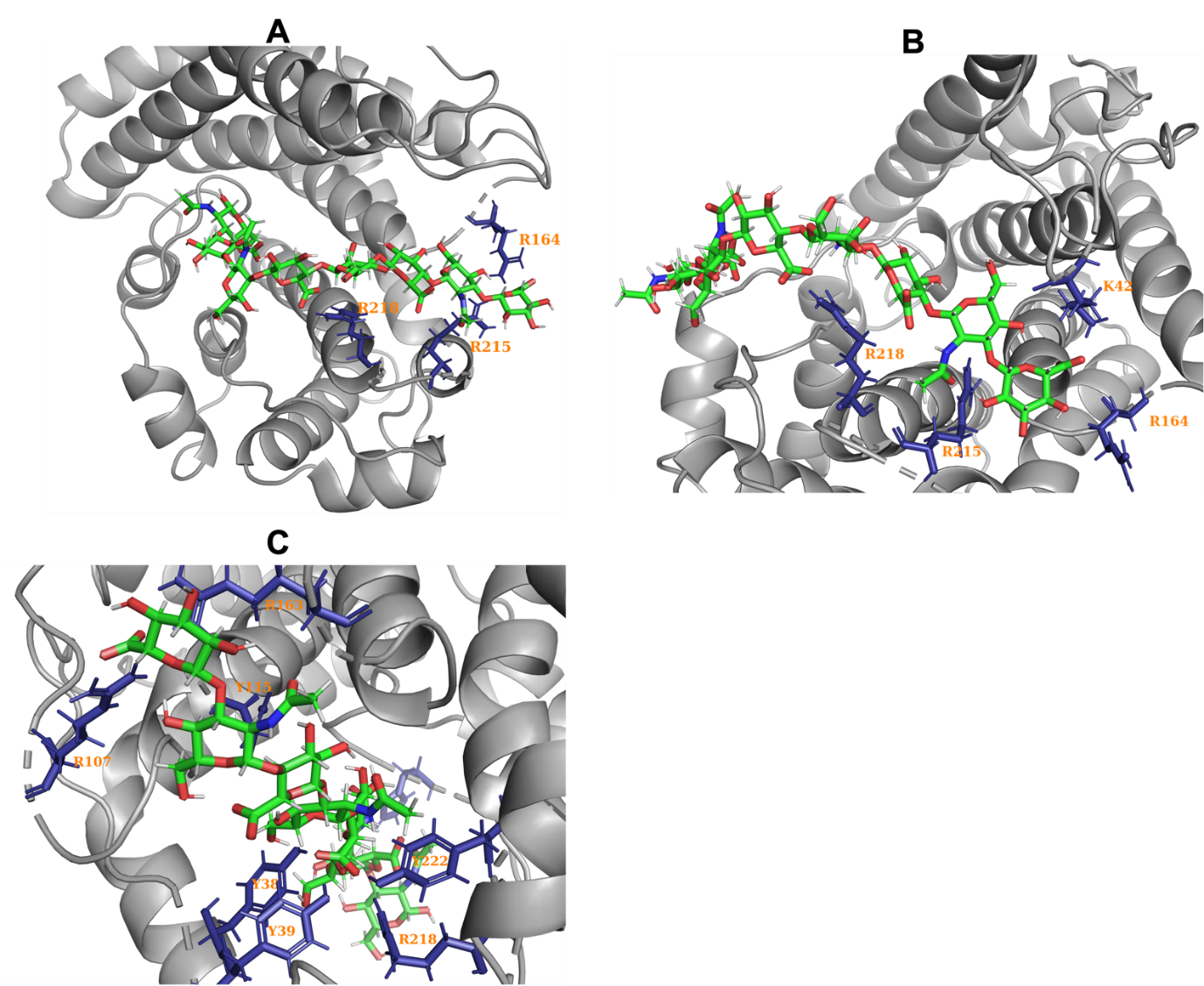

**Figure S18**: The less populated clusters and their major interactions (see text) predicted by our GMVAE algorithm for the WT and H221F simulation with HA_4_. Here are the cluster codes: A- WT cluster 2, B- H221F cluster 0, C- H221F cluster 2.

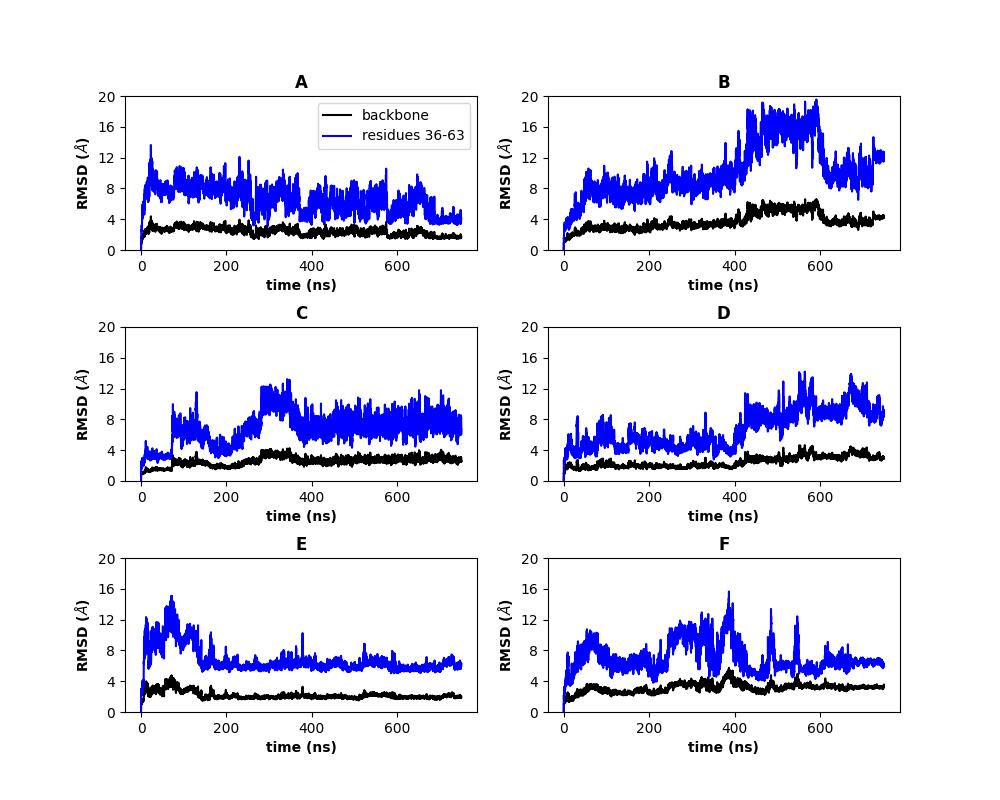

**Figure S19**: RMSD of the protein backbone with $\mathrm{MANA}_{\mathbf{6}}$ as well as the entrance loop with the adjacent helical region (residues 36-63, see Fig. 1B) for A(for WT-set1), B(for WT-set2), C(for H221F-set1), D(for H221F-set2), E(for R312L-set1) and F(for R312L-set 2).

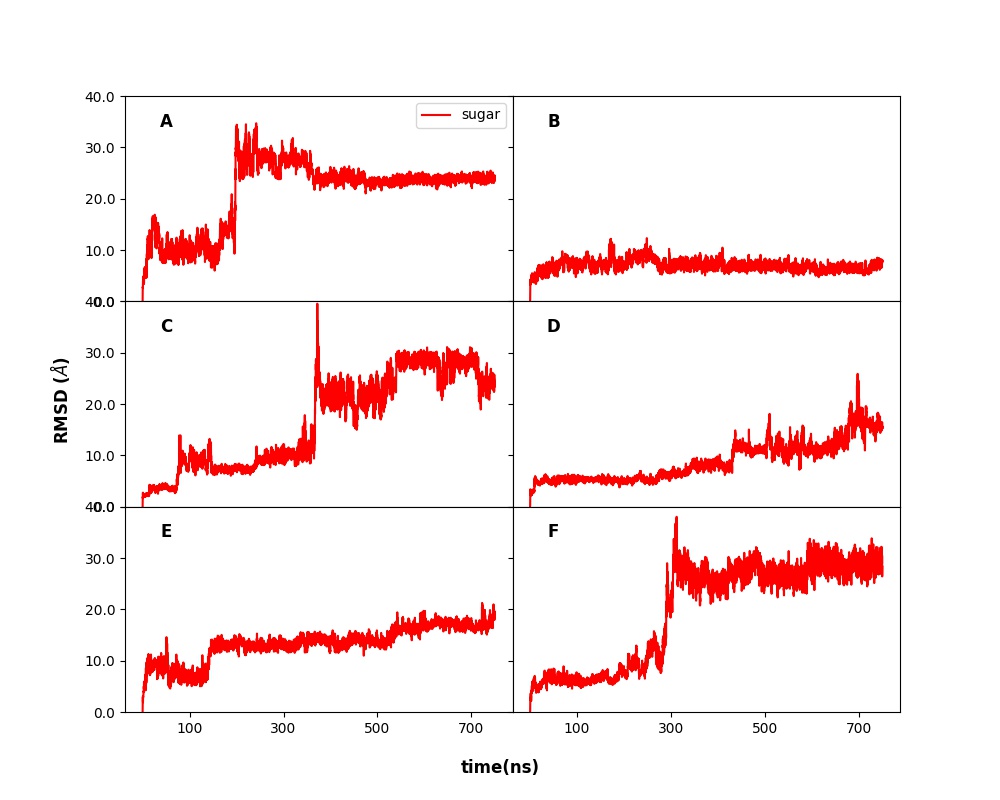

**Figure S20**: RMSD of the sugar $\mathrm{MANA}_{6}$ as a function of time(ns). All these RMSD’s show that the sugar structures oscillate around various regions and that various stable sugar conformations are present inside the binding pocket of the protein for A(for WT-set1), B(for WT-set2), C(for H221F-set1), D(for H221F-set2), E(for R312L-set1) and F(for R312L-set 2).

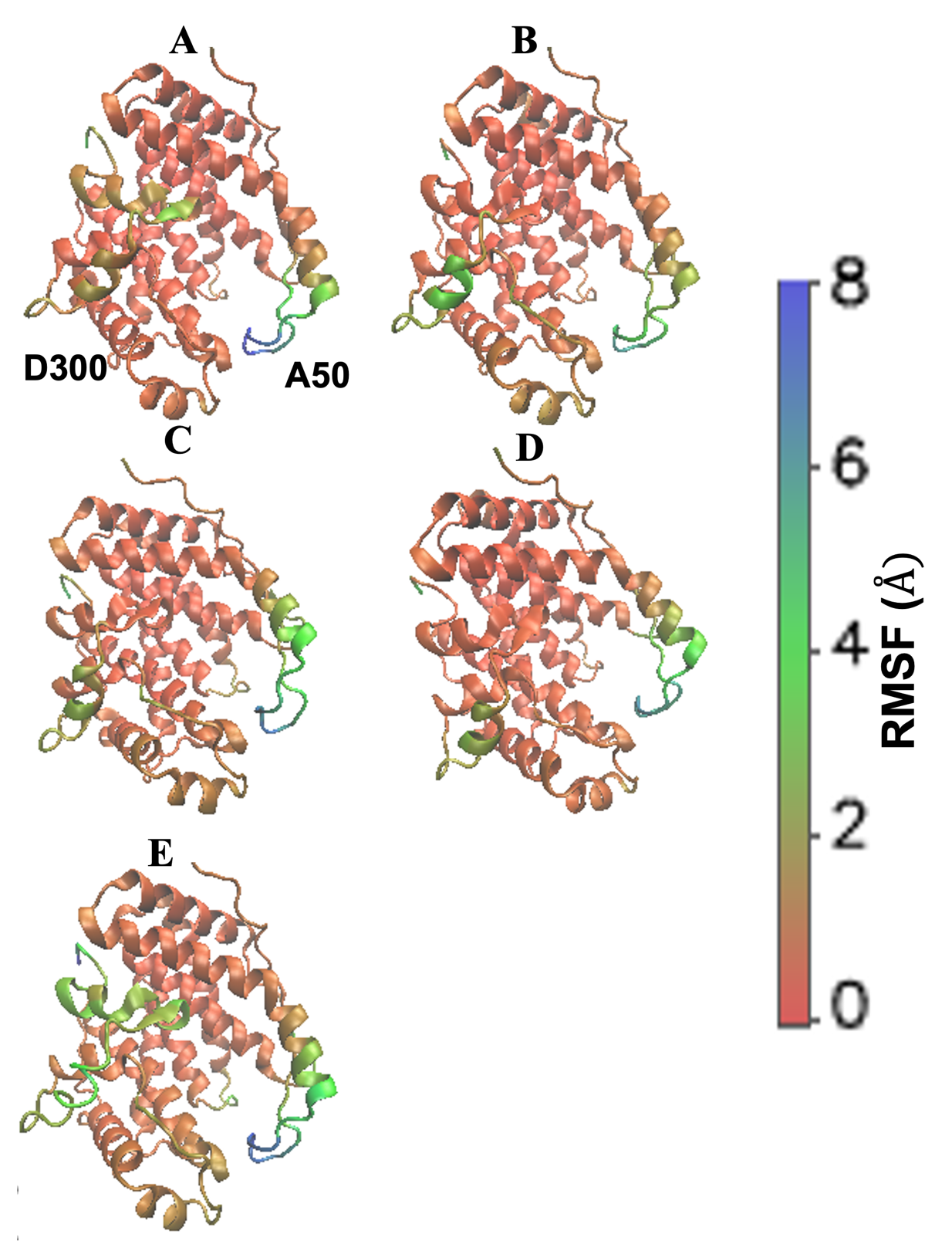

**Figure S21:** RMSF for$\mathrm{MANA}_{6}$ replicas for WT-set2 (A), H221F-set1 (B), H221F-set2 (C), R312L-set1 (D), R312L-set2 (E).

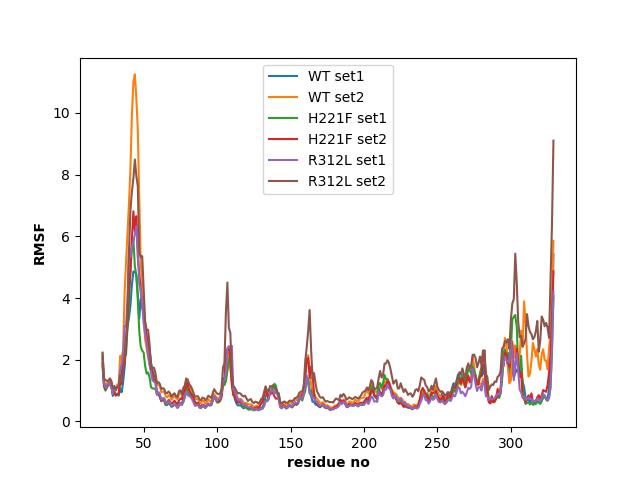

**Figure S22**: The above figure shows the RMSF of the various residues on the protein as a function of the residue no on the protein for our sugar $\mathrm{MANA}_{6}$.

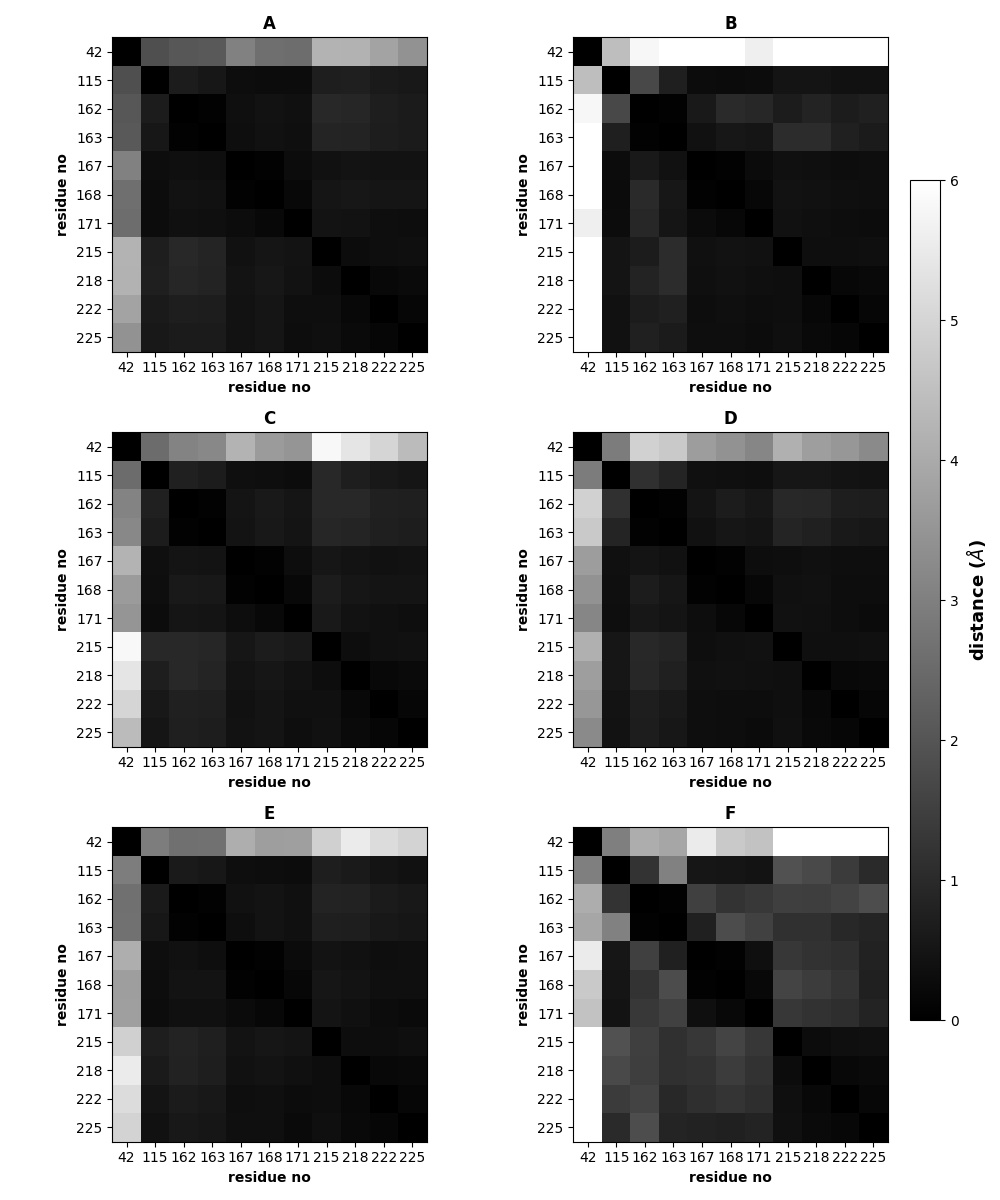

**Figure S23**: A(for WT-set1), B(for WT-set2), C(for H221F-set1), D(for H221F-set2), E(for R312L-set1), F(for R312L-set2) respectively denotes the fluctuations of the distances between various residues on the protein for the sugar $\mathrm{MANA}_{6}$.

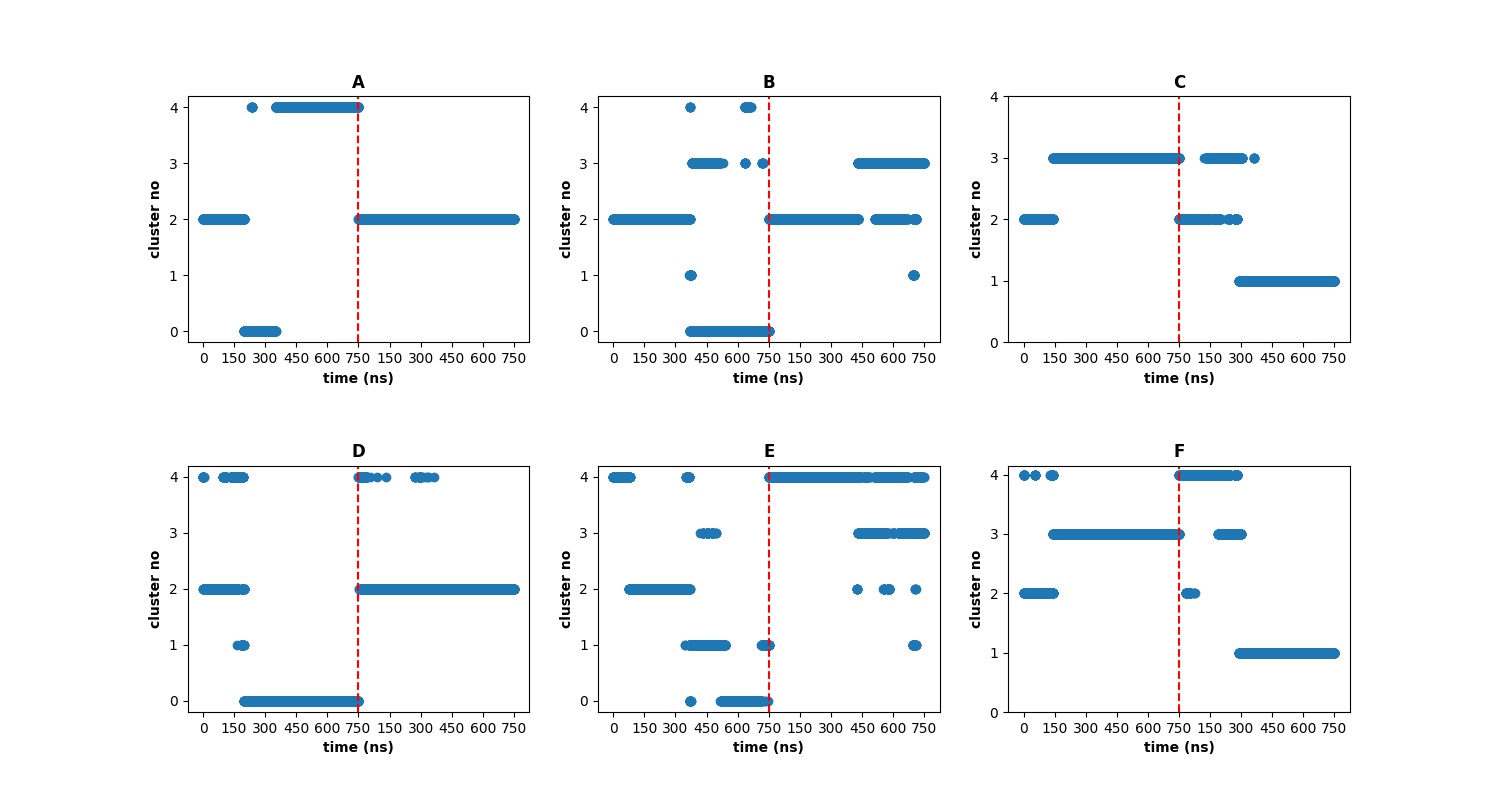

**Figure S24**: A figure showing the various clusters as a function of the simulation time predicted by our KMeans and GMVAE algorithm for both the wild-type, H221F, and R312L mutation. (A), (B), and (C) denotes the results for WT, H221F, and R312L from our KMeans algorithm and (D), (E), and (F) denotes the results from our GMVAE algorithm. We also note the total fraction present for each of the sugar $\mathrm{MANA}_{6}$.

We again note here that on comparing the protein ligand contact maps to the clustering by GMVAE, the GMVAE shows better and closer clustering as compared to KMeans. For example, there is a change in distance for the H221F-set1 in figure S25C after the initial 100 ns which is shown by GMVAE figure S6E but not shown by the KMeans-set B in figure S6B. Similarly for H221F-set1 after 550 ns there is a change in distance which is equivalently reflected through both the protein ligand contact maps as well as the cluster analysis for GMVAE but not through KMeans. For R312L, we see that KMeans show the same cluster 2 for both the initial ranges of the simulations for the two sets, which however is not shown by GMVAE. This variation in cluster by GMVAE is in fact agreement with the protein ligand contact maps which show that there is a disagreement in the colour plots after the initial ranges of the simulation.

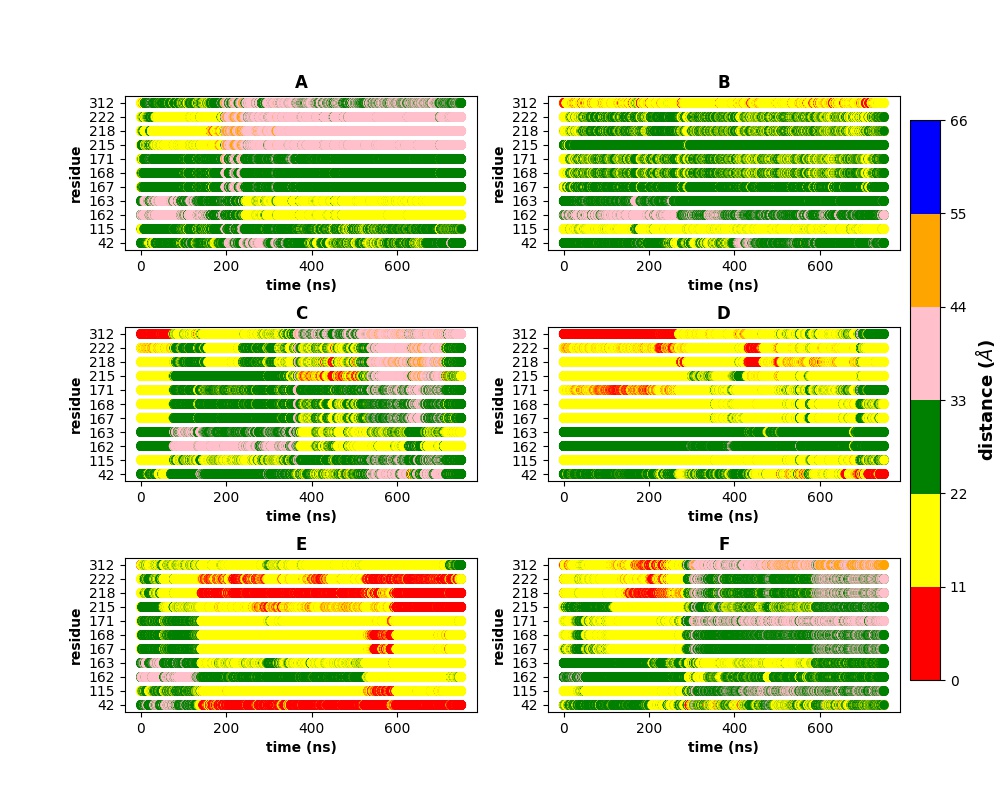

**Figure S25**: A figure showing the various protein ligand distances at various times of the simulation. A(for WT-set1), B(for WT-set2), C(for H221F-set1), D(for H221F-set2), E(for R312L-set1), F(for R312L-set2) respectively denotes the results for the various proteins and the mutants for the sugar $\mathrm{MANA}_{6}$.

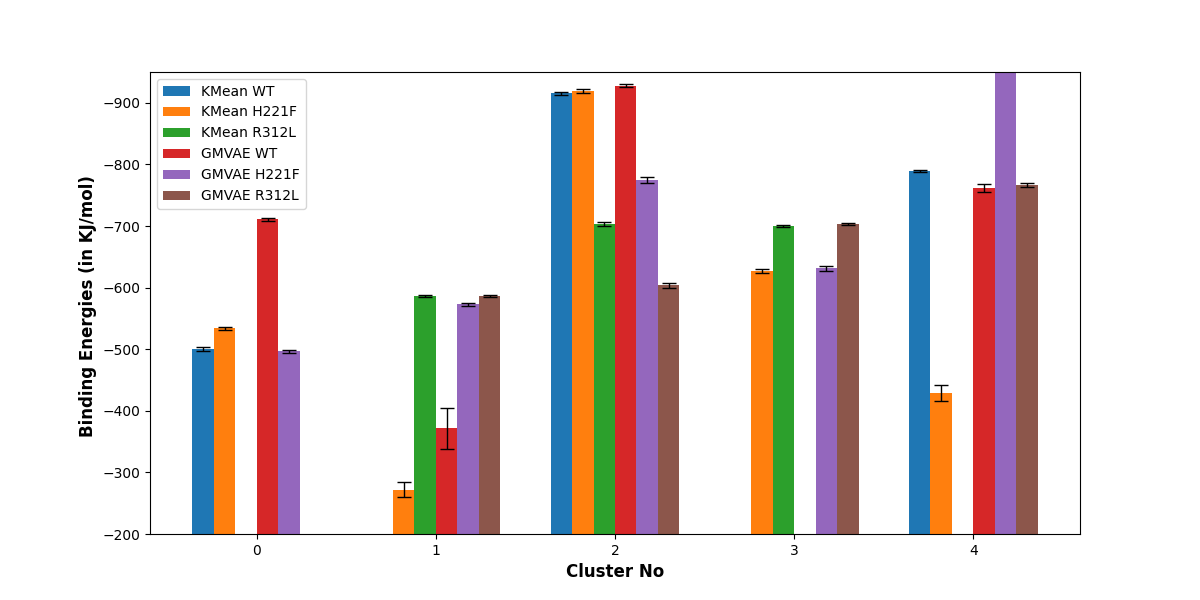

**Figure S26**: The various protein-MANA_6_ binding energies are shown for the various clusters predicted by our KMeans and GMVAE algorithm. The detailed data for this figure is shown in Table S10.

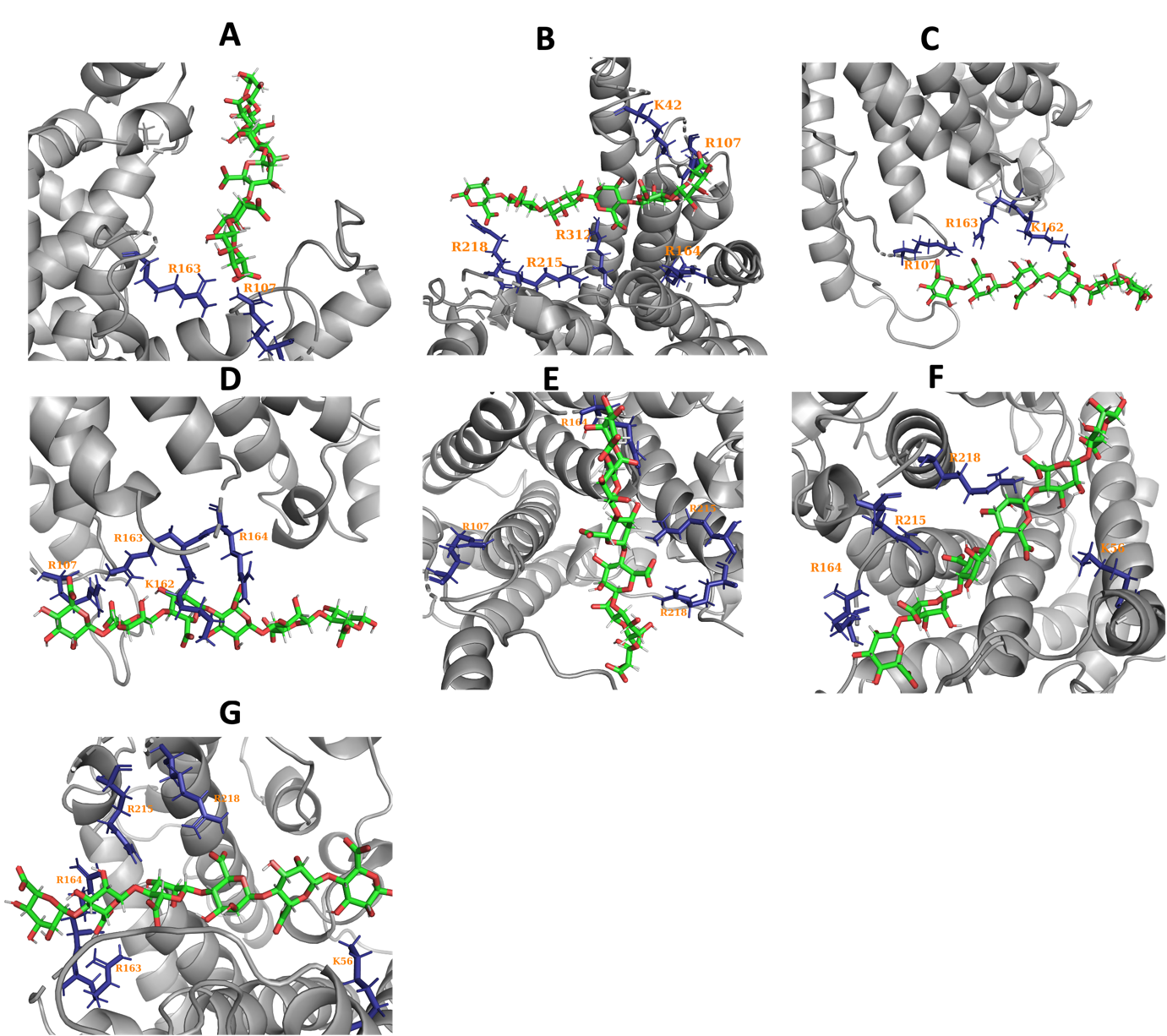

**Figure S27**: The highest 2 populated clusters with the respective highly interacting residues to the sugar from WT simulation ((A) and (B)), H221F mutant ((C) and (D)), and R312L mutant (E), (F), and (G). Here are the cluster codes: (A)- cluster 1 WT, (B)- cluster 4 WT, (C)- cluster 0 H221F, (D)- cluster 1 H221F, (E)- cluster 3 H221F, (F)- cluster 2 R312L, (G)- cluster 4 R312L.
